# A granular view of the traits controlling soil bacterial community reconstruction following a prescribed burn

**DOI:** 10.1101/2024.06.20.599938

**Authors:** Siddharth Uppal, Jamie Woolet, Muthusubramanian Venkateshwaran, Christopher Baxer, Yari Johnson, Ashley Tung, Charlie Siwei Yu, Thea Whitman, Jason C. Kwan

## Abstract

Prescribed burns and wildfires both cause transient reduction of soil microbiome diversity, which has wide-ranging functions in soil and plant health. However, a genetic basis for understanding the mechanisms behind post-fire microbiome recovery is not well-established. Here, we conducted prescribed burns in multiple plots with paired unburned controls, at two prairie locations, sampling each over 5 months. We assembled >300 bacterial genomes that were conserved in both time and space, then identified genomes that were either enriched or depleted post-fire compared to timepoint controls. On a metagenome level, burned and unburned samples were functionally equivalent, but on a species level we determined that post-fire survival is more nuanced than possession of previously hypothesized pyrophilous traits. Our study therefore advances the understanding of how both function and taxonomy contribute to success during the reconstruction of a complex soil microbiome.

## Introduction

Soil microbial communities play critical roles in supporting plant communities, ecosystem productivity^1,2^, and driving biogeochemical cycles^3^, and are amongst the most diverse on Earth^4,5^. Furthermore, most bacteria in the soil remain uncultured, are largely uncharacterized, and many belong to novel taxonomic groups. Capturing all the diversity in soil using sequencing is challenging - perhaps impossible^6^. In principle, more diversity can be captured by increasing sequencing depth; however, such efforts result in diminishing returns, progressively requiring orders of magnitude more sequencing to capture each extra percentage of coverage.

Prescribed burns, which are carried out in order to reduce the risk of uncontrolled wildfires^7,8^ or for ecosystem restoration or management^9^, offer a way to study the microbial ecology of soil in a controlled and reproducible manner. Additionally, since prescribed burns are carried out in service of ecosystem management, it is of inherent interest how prescribed burns affect soil microbiomes and, hence, soil health. Fires typically reduce microbial diversity and biomass in the soil^10–14^, which then gradually returns to pre-fire levels. Therefore, not only are post-fire microbiomes potentially simpler to characterize, but successive timepoints and non-burned controls provide an opportunity to observe the enrichment of specific species after burns. Several genetic traits have been implicated in microbial fire-response through changes in microbiome composition detected through 16S rRNA amplicon sequencing^14–18^, such as ability to form spores, ability to degrade pyrogenic organic matter or fast growth potential to take advantage of the reduced competition immediately after fire. However, there remains a need to more directly connect genetic traits with success post-fire through both genome-resolved methods and controlled assessment of different timepoints.

Here, we analyzed post-burn samples from two restored prairies on the University of Wisconsin – Platteville campus - Pioneer and Rountree. Each site was divided into six 4 m × 5 m plots, which were further divided into burned and unburned subplots, each 2 m × 5 m. Samples were collected at four time points - immediately after the burn, a week, a month and five months after the burn, - and at three depths sampling the O, A1 and A2 horizons (**Fig. 1**). Unburned plots served as controls for seasonal variation, resulting in a total of 288 samples. Analysis of these samples by 16S rRNA gene amplicon sequencing allowed us to target two representative timepoints/samples from Rountree prairie for extremely deep metagenomic sequencing (155 Gbp and 294 Gbp). However, binning of these datasets resulted in just a few high-quality metagenome-assembled genomes (MAGs). Following this, we decided to pivot our approach and sequence more samples at a lower depth, rather than sequencing a few at a very high depth, in order to capture the most diversity. This approach allowed for the stochastic assembly of more high-quality MAGs, which we also showed were present across samples.

**Fig. 1.**
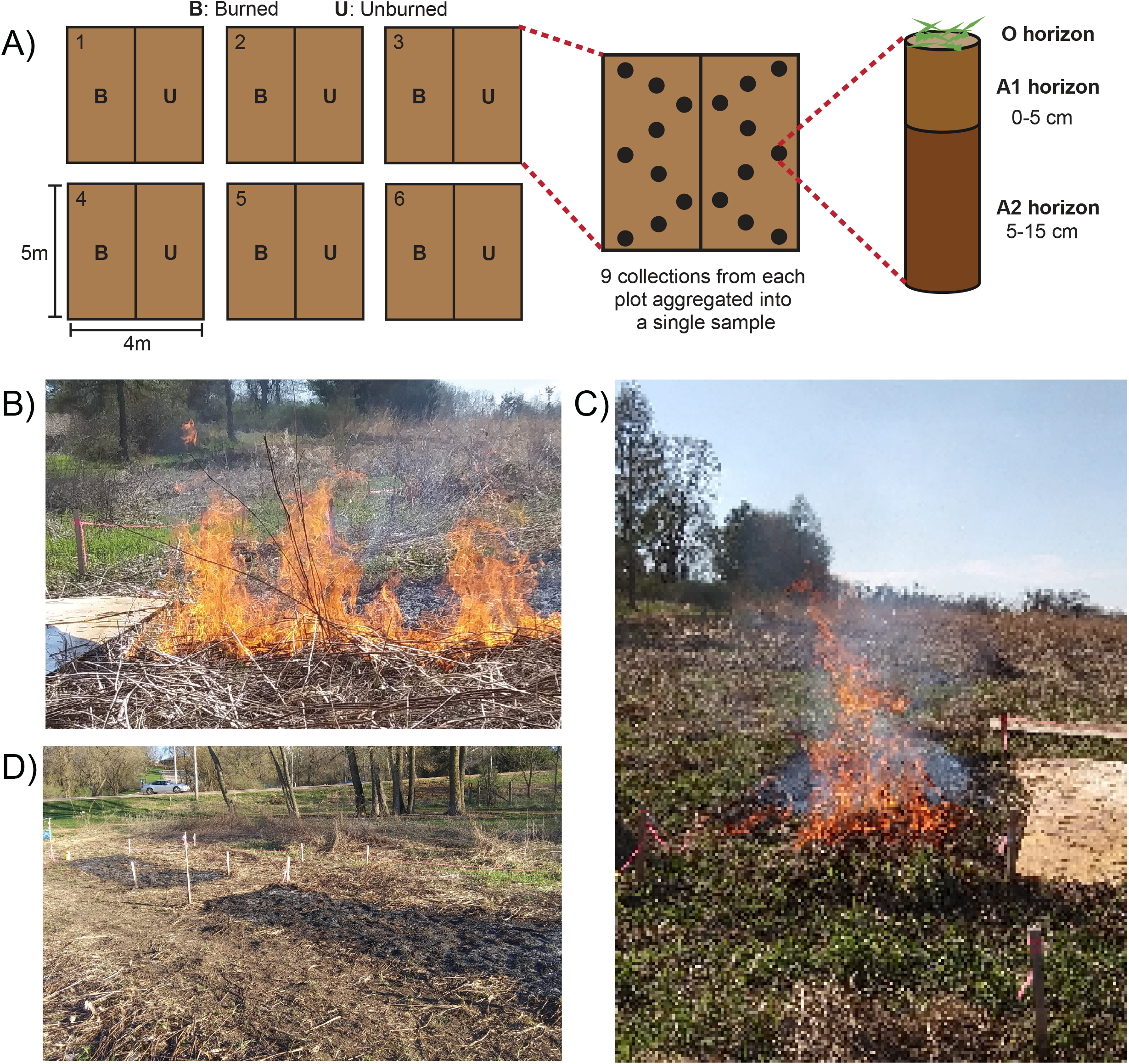
Sample collection and burn site overview. A) Samples were collected from burned and unburned plots at three different depths - and metagenomic data from the top organic (O) horizon was analyzed in this study. Samples were collected at four different timepoints - immediately after, one week, one month, and five months after the burn. B and C) Rountree prairie during the burn. D) Rountree prairie after the burn.

We found that samples collected one week and one month after the burn had distinct microbial compositions between the burned and unburned plots for both the prairies, while communities in burned plots had effectively recovered to match those of unburned communities five months post-burn for the Rountree prairie. Despite significant differences in terms of richness and species composition during the study, metagenomic sequencing revealed that similar functional composition was maintained between the burned and unburned samples throughout, providing evidence for functional redundancy in the soil microbiome and the likely maintenance of functional potential. Drawing on the genome-resolved metagenomic data, we highlight the potential role of spore-related genes and benzoate degradation pathways in supporting increased post-fire abundance of specific taxa, while simultaneously failing to find evidence for any significant role of predicted doubling time and a number of putative post-fire adaptive genes, including those for oxidative stress tolerance and osmotic stress. Critically, we emphasize that having specific fire-responsive genes does not guarantee an increase in relative abundance post-fire. Additionally, we illustrate apparent functional differences between closely related MAGs that might contribute to differential changes in relative abundance post-fire. In such a way, we highlight the complexities of soil microbiomes that can only be revealed through genome-resolved methodology.

## Results and Discussion

### Controlled burns most affected microbiome composition in the O horizon

16S rRNA gene sequencing revealed the greatest difference in microbial diversity between burned and unburned plots in the topmost horizon of the soil (O), compared to lower horizons (A1 and A2) (**Fig 2A B and Extended Data Fig. 1**). The Shannon diversity in the O horizon was significantly reduced one week after the burn, compared to unburned plots. Diversity in the O horizon started to return to unburned levels by a month after the burn and was similar to unburned counterparts five months after the burn (**Fig 2A B)**. However, these trends in reduced and then recovering Shannon diversity were not reflected in the underlying mineral horizons (A1 and A2), indicating that the lower horizons are relatively protected from the impact of the fire (**Extended Data Fig. 1)**, similar to other recent studies of microbial communities impacted by forest fire^10^.

**Fig. 2.**
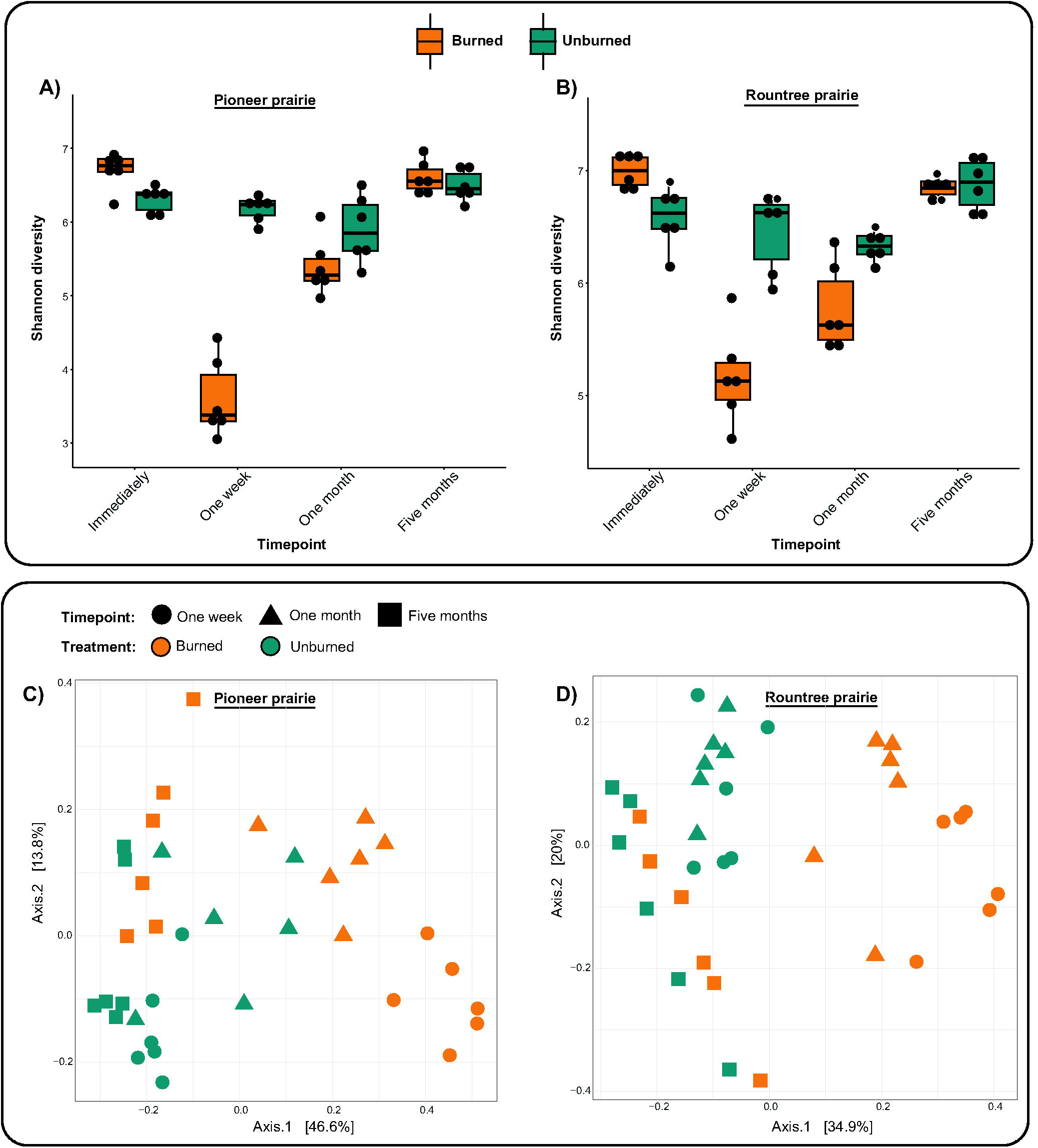
Shannon diversity in the O horizon for A) Pioneer prairie and B) Rountree prairie. PCoA ordination of Bray Curtis dissimilarities for OTU relative abundances in the O horizon for C) Pioneer prairie and D) Rountree prairie.

The effect of fire on the O horizon was further reflected in the community composition of the plots (**Fig 2C D**). In the O horizon, the microbial communities in burned plots differed significantly from the corresponding communities in the unburned plots a week and a month after the burn (PERMANOVA, p < 0.05). In the case of Rountree prairie, this difference was not detectable five months post-burn when the communities started to return to resemble the unburned communities (PERMANOVA, p = 0.123). However, the burned and unburned microbial communities in the Pioneer prairie were still significantly different (PERMANOVA, p = 0.015) from each other five months post-burn, even though the burns at the two prairies were qualitatively similar. This difference between Rountree and Pioneer prairie five months post-burn illustrates that the exact trajectory of recovery of a soil community after a disturbance will be unique to a given system, and could be affected by many different factors. For example, the two prairies differ in slope, and would be expected to have corresponding differences in temperature and moisture (see Materials and Methods for site descriptions). Contrary to these differences in the O horizon communities, the mineral horizons showed no significant differences (PERMANOVA, p > 0.05) between burned and unburned communities at any time point (**Extended Data Fig. 2**).

## Metagenomic analysis

### Metagenome-level gene function profile altered by timepoint but not by burns

Community composition and Shannon diversity for samples collected one week and five months after the burn for Rountree prairie appeared to recover to baseline during the timescale of the study (**Fig 2A C**). This prompted us to sequence the burned and unburned samples of Rountree prairie a week and five months post-burn. For full details on how sequencing depth was chosen refer for **Supplementary Note 1**.

Metagenomic assemblies obtained from Rountree prairie - one week and five months post-burn - were characterized functionally after gene-calling using relative abundance of clusters of orthologous group (COG) counts to get an overview of the functional changes in the soil community post-burn. COG counts were calculated by adding up the number of genes in a sample annotated with a specific COG category. Even though no significant differences (PERMANOVA, p > 0.05) were observed between the burned and unburned plots at any timepoint (**Extended Data Fig. 3AB**), samples from one week and five months formed distinct clusters (PERMANOVA, p = 0.016) (**Fig 3A**). This was in contrast to the results obtained from the clustering of microbial relative abundance data from 16S rRNA gene sequences and MAGs (see below), where clustering based on treatment (burned vs. unburned) as opposed to timepoints was observed. The clusters based on timepoints rather than treatment were apparent when using relative abundance of COG counts, indicating that even though the microbial composition of the soil is altered a week after the burn (**Fig 2**), the potential functional composition remains relatively stable. This is consistent with previous studies that indicate a high level of functional redundancy in many microbial ecosystems^19,20^ and the extremely high diversity typical of soil environments^4,5^. Critically, these findings may be of interest to land managers, as they underscore the notion that even apparently large reductions in measured diversity are not likely accompanied by large functional changes following disturbances such as low-severity prescribed fires.

**Fig. 3.**
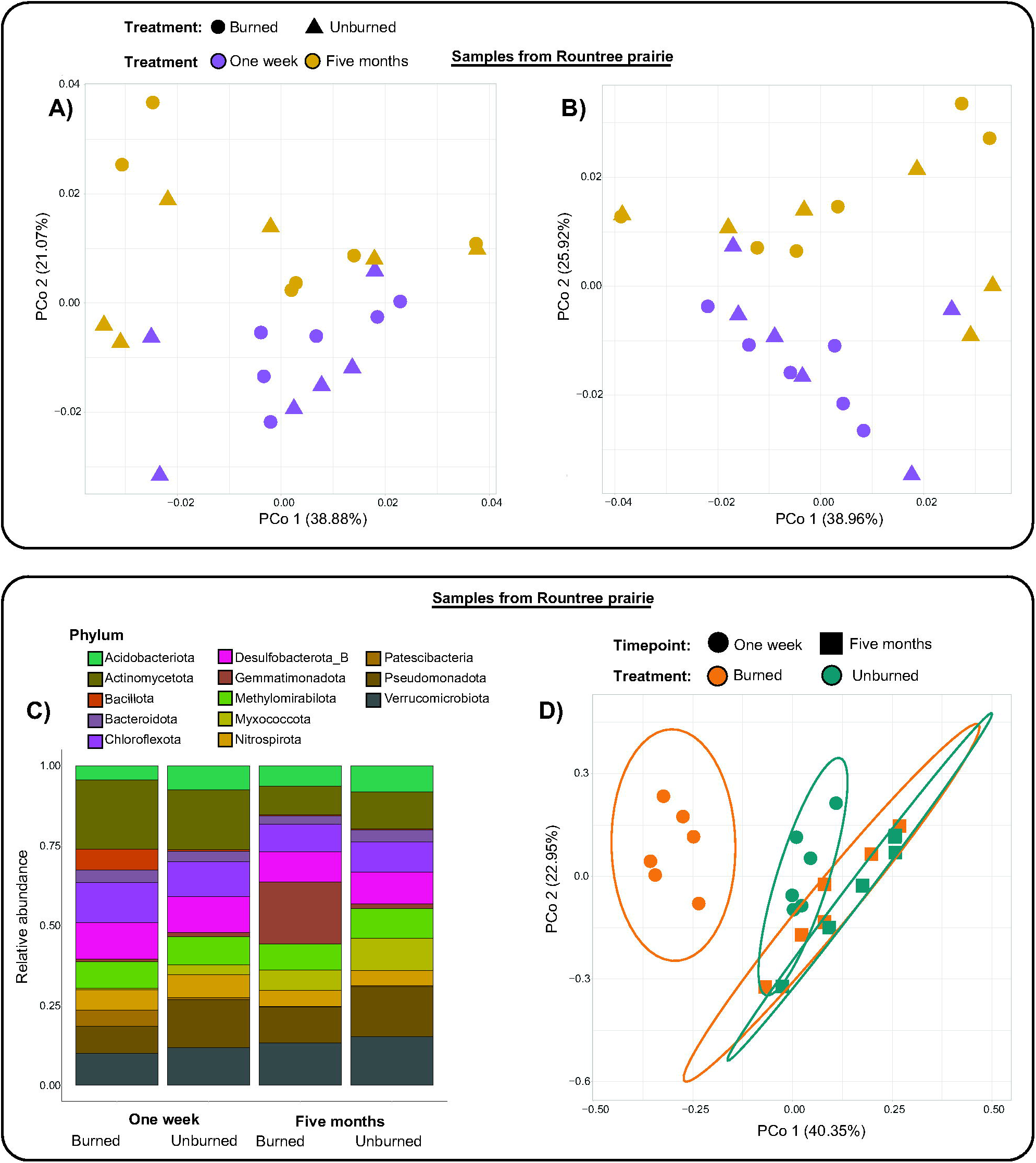
PCoA (using Bray-Curtis dissimilarities) of A) relative abundance of COG counts and B) relative abundance of COG coverage. C) Relative abundance of MAGs of different phyla in four different conditions. D) PCoA (using Bray-Curtis dissimilarities) of relative abundance of MAGs in each sample.

Analysis was also conducted using COG coverage, which was calculated by summing the coverage of the contigs on which the gene annotated with a specific COG category was present in a sample. If multiple genes annotated with the same COG category were present on the same contig, the coverage was added that many times. Clustering based on relative abundance of COG coverage also revealed distinct clusters between samples from a week and five months post-burn (PERMANOVA, p = 0.005) (**Fig 3B**). However, marginal significant differences were also observed between burned and unburned samples a week post-burn - p-value was oscillating between 0.04 and 0.06 using PERMANOVA (**Extended Data Fig. 3CD**). This difference in COG coverage of burned and unburned samples a week post-burn (if statistically significant) might be due to the increase in coverage of fast growing microbes that increase significantly in relative abundance shortly after a burn^21–23^.

### Diverse metagenome assembled genomes (MAGs) conserved in both time and space

Soil has a large diversity of rare and less relatively abundant bacteria^24^, which adds further complexity to the metagenome assembly and binning process, often resulting in few high or medium-quality MAGs that are conserved between samples. To calculate the abundance of MAGs across each of the 24 samples, we first assembled and binned the reads from each sample separately, followed by dereplication of all MAGs together. This resulted in 308 high- and medium-quality bacterial MAGs, as per MIMAG standards^25^. To calculate the relative abundance of each MAG across all 24 samples (**Supplementary Table 1A)**, we aligned reads from all the samples on all 308 MAGs. A global analysis revealed that reads from multiple samples aligned to the majority of the length of MAGs, while there was consistently low alignment to control genomes not expected to be present in soil (**Supplementary Table 1B**). This result indicates that our MAGs are representative of abundant species that are consistently present across sampling sites and timepoints. In some cases, variability in relative abundance of a MAG across different plots was observed. This variability is, however, expected, given the biological diversity of soil. Length weighted coverage was calculated for each MAG in each sample, which was then used to find the relative abundance of that MAG in the sample (**Supplementary Table 1C**). Mean relative abundances of MAGs from the six subplots at each condition (one week - burned/ unburned, five months - burned/unburned) was used to calculate the mean relative abundance of the MAG for that condition. Standard deviation and coefficient of variance for the relative abundance of MAGs across plots have been noted in **Supplementary Table 1D**. MAGs from 13 different phyla including Acidobacteriota (7 MAGs), Actinomycetota (formerly Actinobacteriota) (136 MAGs), Bacteroidota (9 MAGs), Bacillota (formerly Firmicutes) (4 MAGs), Gemmatimonadota (1 MAGs), Patescibacteria (3 MAGs), and Pseudomonadota (formerly Proteobacteria) (128 MAGs) were recovered (**Fig. 3C**).

### Enrichment of both metabolically competent and reduced MAGs post-fire

To identify if a MAG was increasing/decreasing (i.e., enriched/depleted) in burned vs. unburned plots, percentage change in relative abundance was calculated for that MAG relative to the unburned plots at the same timepoint. The metagenomic results discussed below were also corroborated by the differential abundance analysis using the 16S rRNA gene data. MAGs in burned plots a week later were significantly different from their unburned counterparts as opposed to five months after the burn (**Fig. 3D and Extended Data Fig. 4**), further supporting the conclusions from the 16S data. Recapitulating previous results, we identified MAGs related to taxa which are known to be fire responsive across diverse ecosystems including those belonging to genus *Arthrobacter* (n=5, including genera *Arthrobacter_F* and *Arthrobacter_H* as classified by GTDB-Tk)^10,14,15,18,26,27^, *Blastococcus* (n=1)^10,15,23,26^ and *Telluria* (n=2, formerly *Massilia*^28^)^14,15,18,23,26^. We also identified other previously reported fire responsive taxa including *Aeromicrobium*^15,23^ and *Pseudomonas_E*^17^, but not all MAGs belonging to those genera were found to be enriched in burned vs. unburned (**Fig 5**). Our findings add an even more divergent ecosystem (grassland) and fire regime (low-severity prescribed fire) to the growing list of systems in which these taxa have emerged as pyrophilous, while the genetic information in the MAGs informs our emerging understanding of bacterial fire ecology.

**Fig. 4.**
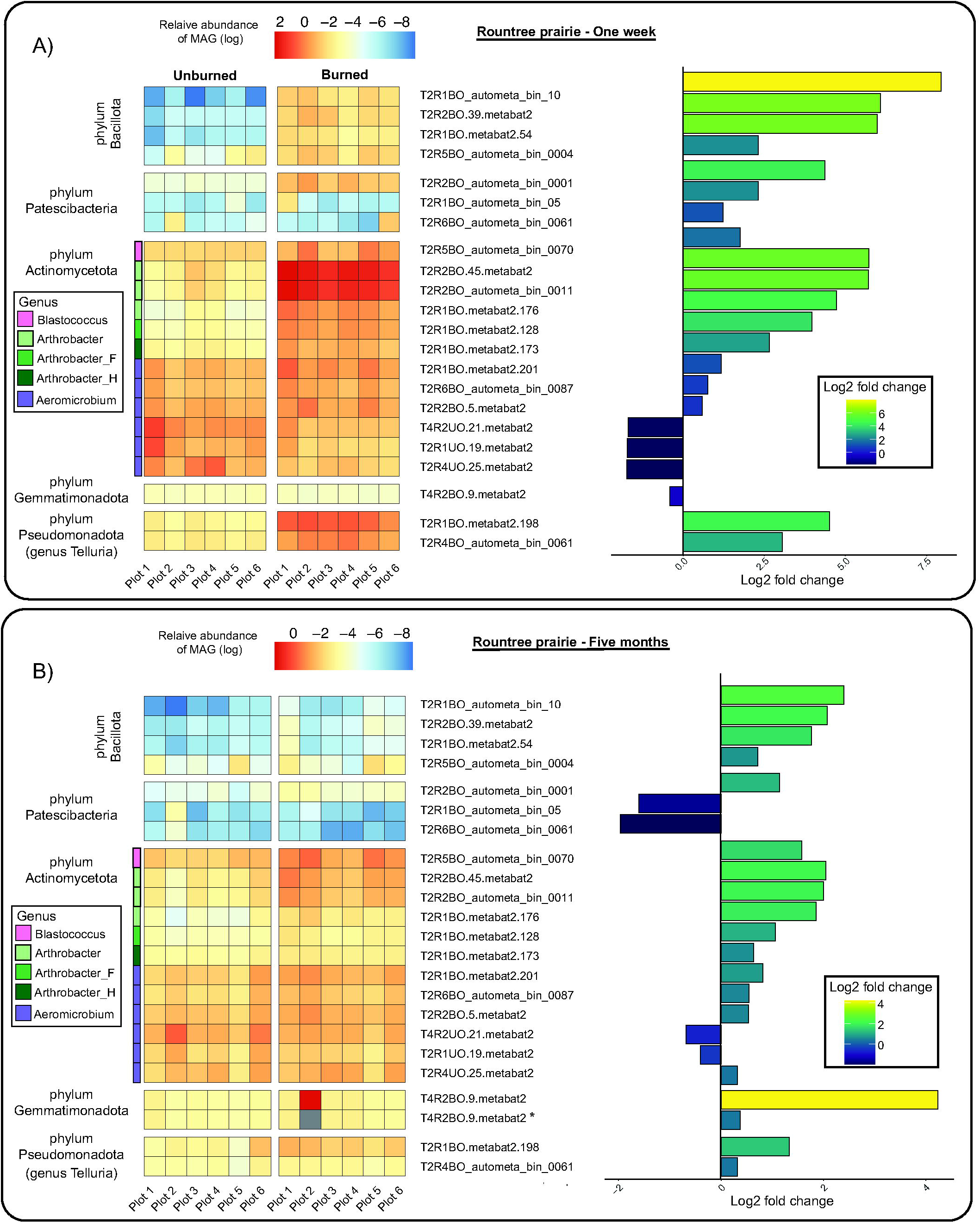
Heatmap and log2-fold change for selected MAGs from rountree prairie. A) A week after burn, B) Five months after burn. *Log2-fold change calculated for MAG belonging to phylum Gemmatimonadota (T4R2BO.9.metabat2) without taking sample T4R2BO (grayed out) into account.

**Fig. 5.**
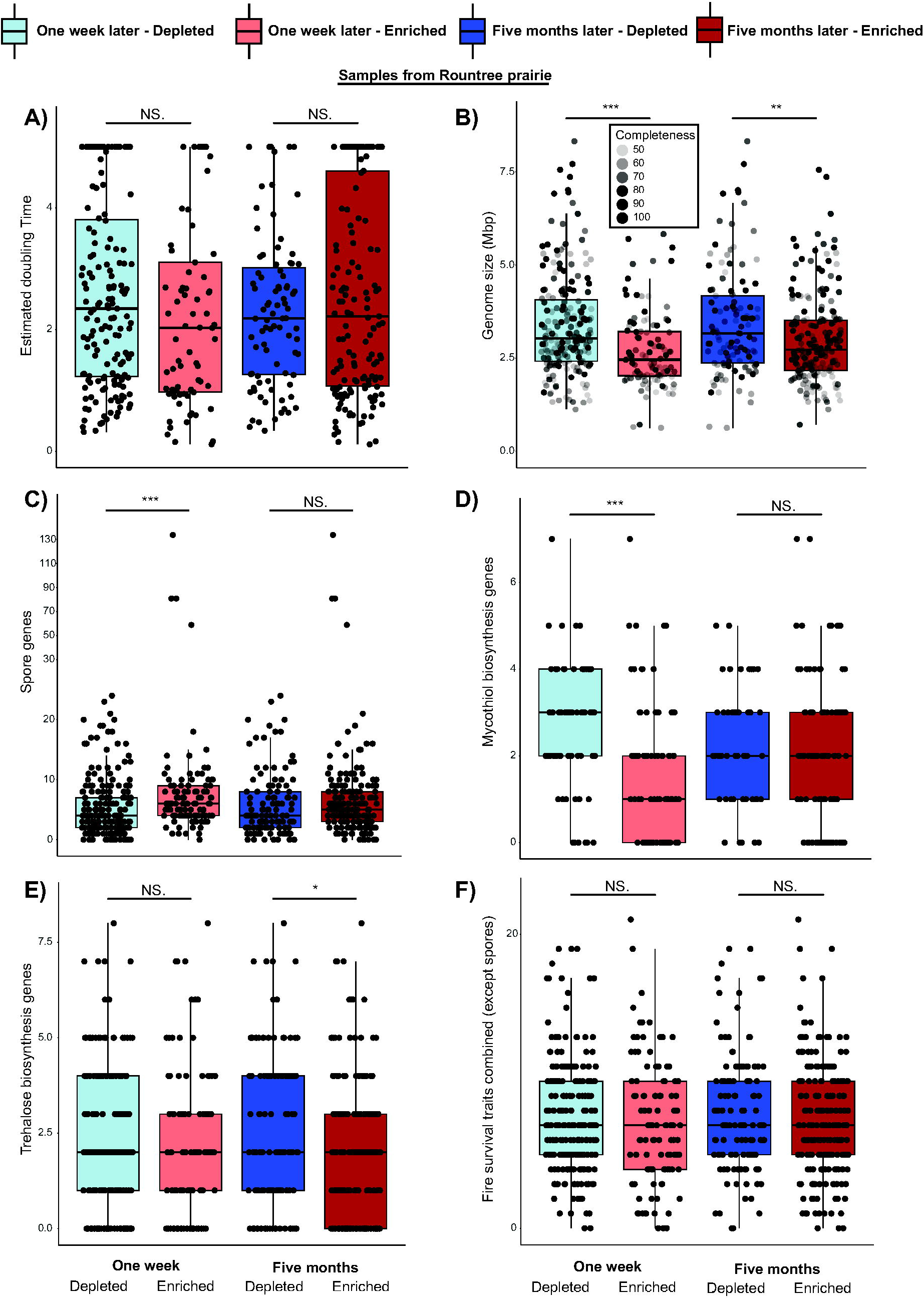
Characteristics for all MAGs classified by whether they are depleted or enriched in burned vs. unburned plots one week and five months post-fire from rountree prairie. A) Estimated doubling time. Doubling time of 5h is used for MAGs when estimated doubling time is > 5h (See methods for full details). B) Genome size. C) Spore gene count; note scale on y-axis is reduced by a factor of 5 after 30. Number of genes for D) mycothiol biosynthesis and E) trehalose biosynthesis. F) Gene count for all fire survival traits (except spore genes) combined; note that scale on y-axis is reduced by a factor of 5 after 60. NS indicates no significant difference between the groups. * indicates significance at p < 0.05, ** indicates significance at p < 0.01, and *** indicates significance at p < 0.001.

Surprisingly, MAGs belonging to phylum Patescibacteria (n=3) (a single phylum formerly classified as the candidate phyla radiation)^29^, which have extremely small genomes and lack genes for numerous biosynthetic pathways^30,31^, had one of the largest increases in relative abundance in burned vs. unburned plots one week post-burn. Patescibacteria have been observed to survive as symbionts with other bacteria^32–35^, eukarya^36^ and even archaea^37–40^. Patescibacteria MAGs recovered in this study belonged to class “*Candidatus* Saccharimonadia” (formerly known as TM7), whose members have been reported to have an epibiotic parasitic (or epiparasitic) lifestyle with different Actinomycetota including members of genus *Rhodococcus*, and *Nocardia*, amongst others^32,35^. Such parasitic relationships could explain the increased abundance of MAGs with reduced metabolic capabilities. Our metagenomic analysis did identify MAGs belonging to *Rhodococcus*, and the relative abundance of one of them (T2R4BO_autometa_bin_0042) was co-abundant with one of the Patescibacteria MAGs (T2R2BO_autometa_bin_0001) post-burn, but we do not view this as sufficient evidence to establish a parasitic relationship. Further research would be warranted to identify and verify the presence of a parasitic association of Patescibacteria with *Rhodococcus* in soil.

Two of the “*Ca.* Saccharimonadia” MAGs, belonging to family UBA4665 (T2R6BO_autometa_bin_0061 and T2R1BO_autometa_bin_05), were enriched in burned vs. unburned samples a week post-burn but were depleted five months later. In contrast, the “*Ca.* Saccharimonadia” MAG belonging to family UBA1547, genus *UBA6022* (T2R2BO_autometa_bin_0001) was enriched in burned vs. unburned at both timepoints. The fact that the Patescibacteria were found to be enriched in burned vs. unburned plots post-fire indicates that they may have heat or stress-tolerant adaptations, thrive in a post-fire environment and/or have hosts adapted to post-fire environments. However, the increase of only one of the three MAGs five months later could indicate that the initial opportunistic growth of Patescibacteria may have been dependent on specific post-fire conditions (e.g., fire-liberated nutrients from fire-killed organisms or fire-altered organic matter) that diminished over time. Though it is hard to confidently discern any symbiotic associations with sequencing data alone, these results open the door to answering how metabolically deficient bacteria are able to opportunistically survive and even thrive post-fire. Furthermore, the fact that Patescibacteria MAGs were found to be present after a wildfire by another recently published study^10^ underlines the need to study the post-fire survival adaptive traits of this phylum in further detail.

A MAG (T4R2BO.9.metabat2) belonging to phylum Gemmatimonadota (family Longimicrobiaceae, genus *JAFAYN01*), was the most enriched in burned vs. unburned plots, out of all the MAGs, five months after the burn (**Fig. 4**). However, it is important to note that T4R2BO.9.metabat2 had an exceptionally high relative abundance in the sample T4R2BO compared to all the other plots (69.08× compared to the plot with second highest relative abundance), likely because that is where it was originally binned from. After removing sample T4R2BO from the analysis, T4R2BO.9.metabat2 was still enriched in burned vs. unburned five months post-burn, although the log2 fold increase was reduced from 4.24 to 0.37 (**Fig. 4**). Genomes belonging to phylum Gemmatimonadota have been recently found to be enriched after a prescribed burn^41^ and have also been shown to respond positively to PyOM additions^42^. However, inspection of the MAG revealed the presence of very few putative PyOM degradation genes and no complete set of genes for any aromatic degradation pathway inspected in this study. Thus, initial genetic evidence for this MAG does not support the hypothesis that they are likely directly degrading aromatic PyOM.

We found that the Gemmatimonadota MAG in this dataset has 153 biosynthetic gene clusters (BGCs) predicted to make secondary metabolites, mostly non-ribosomal peptide synthetases. Since the MAG is fragmented, the BGC count could be artificially high, with the same BGC being fragmented on multiple contigs. However, we found that the BGCs made up 1.20 Mbp or 16.36% of the entire genome when calculated using the length of the proto core region, i.e., the core biosynthetic genes, and 1.69 Mbp or 23.02% of the entire genome when calculated using the length of the protocluster region of the BGC, which includes the core genes and genes encoded up- and downstream of the core (as determined by antiSMASH rules^43^). Soil microbes have been shown to be rich in BGCs that produce secondary metabolites^44–46^. Furthermore, a recent study of 3304 soil metagenomes also highlighted the rich biosynthetic potential of Gemmatimonadota^46^. We initially hypothesized the increased coding capacity for BGCs in the Gemmatimonadota MAG will confer it a selective advantage post-burn, due to its ability to synthesize natural products and reduce competition. However, even though the MAG appears to be quite enriched in burned vs unburned five months later, BGCs don’t appear to offer a clear advantage in this MAG across all or even most burned plots five months later.

### Fire response traits

#### Post-burn microbiome enriched in smaller, high-GC MAGs

Recolonization of the soil by copiotrophic bacteria - characterized by fast growth and the ability to thrive in nutrient-rich environments - is expected to be an important post-fire disturbance response^15^. We used gRodon2^47^, an R package, to predict minimal doubling time using genome-wide codon usage statistics. We expected that the fire-induced death of bacteria, fungi and plants may have created a nutrient rich environment favorable for copiotrophic bacteria to thrive in, making copiotrophic characteristics in MAGs an important trait for post-fire success. Although we observed a slightly lower median of predicted doubling time for MAGs enriched in burned vs. unburned one week post-burn, the differences between predicted doubling times in MAGs enriched and depleted in burned vs. unburned were not significantly different at any time point (**Fig. 5A**). The lack of a clear signal of decreased predicted doubling times in MAGs enriched post-burn could reflect a number of possibilities. First, we may not have captured the timescale over which this phenomenon is relevant. In the boreal forest of Canada, the importance of fast growth was present in high severity fires one year post-burn, but not five years later^21^. For a low-severity prescribed fire in a more temperate climate, where recovery might be expected to be faster, any importance of fast growth may be particularly short-lived. Second, it is important to note that predicted doubling time does not necessarily reflect the actual activity of a given MAG - fast growth may still be an important trait over the timescales we measured here, but this metric does not capture it.

Recent studies have reported small genome size of bacteria from soil to be associated with stress tolerance. Piton *et al*. used the CSR (competitor, stress tolerant, ruderal) trait framework to study the life history strategies of soil bacterial communities and identified bacteria with streamlined genomes to be associated with stress tolerance^48^. They further observed smaller genomes to be associated with arid environments^48^, something that has also been reported in another study^49^. This is in line with the observed enrichment of three Patescibacteria MAGs a week post-burn. Moreover, Simonsen studied the pangenome of soil bacterium *Bradyrhizobium diazoefficiens* across different environments and identified strains isolated from higher stress environments (more acidic, more saline, hotter and drier) to have a smaller genome size^50^. Here, we observed that MAGs that were more relatively abundant in burned vs. unburned a week and five months after burn had a significantly smaller genome size (Wilcoxon test, p = 2.20e^-06^ for week post-burn and p = 0.007 for five months post-burn) compared to those that were less relatively abundant at the same time point (**Fig. 5B**), indicating that streamlined genomes might be more tolerant to fire-associated stress. Since we are dealing with MAGs which can be incomplete, we reanalyzed the above relationship with MAGs that were at least 70% complete, based on checkM2^51^ results, and observed the differences in genome size to be still present one week later (Wilcoxon test, p = 0.035) but not five months post-burn (**Extended Data Fig. 5a**). The lack of significance at five months post-burn might be due to fewer MAGs with completeness greater than 70%. However, further studies using more complete genomes are needed to verify these results. One week after the burn, the percentages of MAGs that were less than 70% complete and below the median genome size in their respective categories (3.04 Mbp for decreasers and 2.45 Mbp for increasers) were about the same (40.09% for decreasers and 43.56% for increasers), suggesting that genome completeness did not bias this finding.

Interestingly, we also identified MAGs enriched in burned vs. unburned one week post-burn to have a significantly higher (Wilcoxon test, p = 0.007) length-weighted guanine-cytosine (GC) content compared to MAGs depleted in the same time point (**Extended Data Fig. 5b**). GC base pairing is known to have a higher thermal stability than adenine-thymine (AT) base pairs^52^. We hypothesized that a genome with a higher GC content could potentially have a survival advantage over one with higher AT content during a fire. In a similar vein, a recent study reported MAGs assembled from surface soil impacted by high severity wildfires to have higher GC contents compared to MAGs from deeper horizons^10^. Johnson *et al*. propose that the importance of fire survival as a trait is likely most important within a relatively narrow range of fire temperature, critically, where there is differential survival^21^. Because our study utilized relatively low-severity fires, it may be the case that the temperatures organisms were exposed to were within the critical range where some taxa could survive, but others could not (rather than being so high that most taxa die, and post-burn composition dynamics are dominated by dispersal and secondary succession).

#### MAGs able to degrade pyrogenic organic matter enriched post-burn

Pyrogenic organic matter (PyOM) is produced by the incomplete combustion of organic matter. It can represent large fractions of total soil organic carbon in ecosystems affected by fire^53^, and is chemically characterized by a relatively high degree of aromaticity compared to non-fire-affected organic matter^54^. The ability of microbes to thrive in a post-fire environment has often been linked to their putative ability to utilize PyOM^15,55^. We investigated the completeness of different aromatic degradation pathways and their effect on post-fire microbial abundance. Amongst the aromatic pathways investigated, we observed that only the MAGs with all the genes for degradation of benzoate^56^ to 3-oxoadipate were consistently enriched in burned vs. unburned after fire at both timepoints (**Extended Data Fig. 6**). Four MAGs met this criteria, two belonging to family Pseudomonadaceae (T2R6UO_autometa_bin_0026 - *Pseudomonas_E mandelii;* and T2R5BO_autometa_bin_0008 - *Pseudomonas_E* sp.), and two belonging to order Burkholderiales, from family Burkholderiaceae_B (T2R1BO.metabat2.154 - *Comamonas* sp.) and family Burkholderiaceae (T4R3BO.57.metabat2 - *Duganella* sp.). Interestingly, none of the five MAGs had all the genes required for the last two steps of benzoate degradation (3-Oxoadipate to 3-Oxoadipyl-CoA by *pcaIJ* and 3-oxoadipyl-CoA to acetyl-CoA and Succinyl-CoA by *pcaF* and *fadA*) (**Extended Data Fig. 6**). The two MAGs belonging to order *Burkholderiales* had *pcaIJ* while the two belonging to family *Pseudomonadaceae* had *pcaF*, while only *Duganella* sp. had the *fadA* gene. This finding hints that exchange of metabolites might be happening, as hypothesized in another recent study^10^. The acetyl-CoA and succinyl-CoA produced at the end of the benzoate degradation pathway can then be used in a number of reactions within the cell, including the citric acid cycle, to increase the amount of energy produced, therefore directly benefiting the survival of the microbe post-fire. Thus, these findings are consistent with the hypothesis that some taxa may be (and remain) enriched post-fire due to their ability to degrade PyOM.

#### Post-fire success not easily predicted from the presence of putative post-fire survival traits

Bacterial spores are known to be highly resistant to extreme environmental conditions, including fire^57,58^. There are a number of different modes of sporulation^59–61^, including endospores - spores formed inside the cell - and exospores - spores formed outside the cell^62–64^. Endospores are believed to be restricted to the members of the phylum Bacillota^65,66^, whereas exospores are commonly found in Actinomycetota genomes. All four MAGs belonging to phylum Bacillota were enriched after the burn. These four MAGs were classified as *Domibacillus tundrae*, *Psychrobacillus psychrotolerans* and *Paenisporosarcina* sp. (n=2). Even though Bacillota are present in relatively low abundance in soil, 3 out of 4 MAGs showed the highest increase in burned vs. unburned a week after the burn and remained amongst the top 10 increasers five months after the burn. This relatively large increase in the relative abundance of Bacillota is often attributed to their ability to form spores and survive high-temperature fire conditions^18,67–69^. We were able to identify a large number of spore genes in the four Bacillota MAGs - 134 in *D. tundrae*, 81 in *P. psychrotolerans* and 81 and 59 in the two *Paenisporosarcina* sp. MAGs - thus providing further evidence supporting the above hypothesis. The total number of spore genes amongst these four MAGs accounted for about 16.92% of all the spore genes identified amongst all 308 MAGs.

The number of spore genes in MAGs enriched in burned vs. unburned one week post-burn were found to be significantly higher (Wilcoxon test, p = 3.13e^-05^) than spore genes in MAGs depleted in the same time point (**Fig. 5C**). This was true even after removing Bacillota MAGs (Wilcoxon test, p = 0.0002), which might be considered outliers with an extremely high number of spore genes compared to the rest of the MAGs (**Extended Data Fig. 5c**). Moreover, not all MAGs with spore genes were enriched in burned vs. unburned a week post-burn. This was especially true for MAGs belonging to Actinomycetota, a phylum whose post fire increase has generally been attributed to their ability to form spores^26,70,71^. We identified that numerous putative spore-forming Actinomycetota were actually reducing in relative abundance a week (64/134) or five months (47/134) after the burn. Furthermore, the median number of spore-related genes in Actinomycetota MAGs enriched post-burn was lower than the median number of spore-related genes in Actinomycetota MAGs that were depleted post-burn (6.5 vs. 7.5) (**Extended Data Fig. 5d**). This discrepancy in post-burn abundance of Actinomycetota and spore genes could be due to a number of factors. Though it might be true that spore genes can protect a microbe from fire and allow it to increase in relative abundance later on, (1) the genes alone are not useful if pre-fire conditions have not led the microbe to actually produce spores, (2) the mode of sporulation (endospore vs. exospore) may affect the post-burn abundance, where exospores are thought to have lower resistance to heat and desiccation compared to endospores^62^, (3) fire survival alone may not be a relevant post-fire trait, particularly if an organism is not able to grow relatively rapidly to take advantage of post-fire reductions in community size. This last trend is also highlighted in the fire traits framework presented by Johnson *et al*.^21^, who found that strong fire- survivors only represented small fractions of the total community one and five years post-fire.

Heat-shock proteins (HSPs) are known to be produced in response to stressful conditions. We initially hypothesized that there might not be a significant difference between in the number of HSP genes between MAGs enriched/depleted in burned vs. unburned as the fire conditions may be so fleeting that microbes are not able to respond to them by expressing HSP-related genes that could maintain cellular functioning at higher temperatures. Although the upper quartile (Q3) of the box plots reported the count of HSPs was higher in MAGs enriched in burned vs. unburned at both timepoints, we only observed a significant difference at five months post-burn, with the p-value barely below 0.05 (Wilcoxon test, p=0.047) (**Extended Data Fig. 7a**). This was surprising but in line with previous studies that have also shown that MAGs increasing after a wildfire tend to have a higher number of heat-shock proteins (HSP)^10^.

Fires alter soil chemistry, often creating oxidative environments, leading to redox stress for microbes^72–76^. Glutathione is the major thiol buffer found in mainly gram negative bacteria^77–79^, which aids in oxidative stress tolerance by acting as a reducing agent within the cell^77^. Actinomycetota lack glutathione; instead, they use another thiol called mycothiol to maintain their cellular redox homeostasis^80^. Furthermore, an enrichment of genes for mycothiol biosynthesis have also been cited as a strategy adopted by MAGs to survive redox stress caused by wildfires^10,73^. While no significant difference in the number of glutathione biosynthetic genes was observed between the MAGs enriched/depleted in burned vs. unburned five months after the burn, we observed a significantly higher (Wilcoxon test, p = 0.035) number of glutathione biosynthesis genes in MAGs decreasing in burned vs. unburned one week after fire compared to MAGs increasing a week post-fire (**Extended Data Fig. 7b**). A similar trend was observed with mycothiol biosynthesis genes, where Actinomycetota MAGs depleted in burned vs. unburned one week post-fire had significantly higher (Wilcoxon test, p = 0.0001) number of mycothiol genes compared to Actinomycetota MAGs enriched post-fire (**Fig. 5D**). These results represent a different picture than previously discussed in the literature, where the presence of mycothiol biosynthesis genes have been correlated with improved post-fire survival^10,73^. One possible explanation for this discrepancy could be that our prescribed burns didn’t reach high enough temperatures to result in soil environments with high redox stress^72,81^.

Previous studies^10,73^ have also reported an increase in genes for the biosynthesis of compounds involved in osmotic stress tolerance post-fire, such as ectoine, glycine betaine and trehalose^82–84^, as an adaptive strategy for post-fire survival. However, we didn’t observe any significant difference in the number of genes for ectoine biosynthesis in the MAGs increasing and decreasing in burned vs. unburned post fire at any time point (**Extended Data Fig. 7c**). We, however, observed that genes for glycine betaine and trehalose biosynthesis were significantly higher (Wilcoxon test, p = 0.029 for glycine betaine biosynthesis and p = 0.011 for trehalose biosynthesis) in MAGs depleted in burned vs. unburned at one week and five months post-burn respectively (**Extended Data Fig. 7d and Fig. 5E**). This is surprising, as genes for trehalose and glycine betaine, both known osmoprotectants^82,83,85^, have previously been reported to be enriched in MAGs abundant one year post-burn^10^.

Because we would predict that numerous traits interact to confer post-fire success, we thought that even though genes for one single trait may not be significantly higher, a combination of these traits may confer post-fire resilience to the microbes. After combing the genes for the traits discussed above - spores, HSPs, glutathione biosynthesis, mycothiol biosynthesis, ectoine biosynthesis, glycine betaine biosynthesis and trehalose biosynthesis - we did indeed observe a significantly higher number of genes in MAGs increasing in burned vs. unburned a week post-fire (Wilcoxon test, p = 0.012) (**Extended Data Fig. 8b**). However, this difference disappeared after removing spore genes (Wilcoxon test, p = 0.579) (**Fig. 5F**). Together, we do not find evidence that genes for oxidative stress tolerance or osmotic stress response provide a distinct survival advantage to MAGs enriched in burned vs. unburned after the prescribed burn. This may reflect the relatively low severity of the prescribed burns in this study - soils did not remain bare over the course of the growing season, as vegetation regrew, likely reducing stressful conditions. It is also possible that some of the genes may have been missed due to the incompleteness of the MAGs or the search criteria used for identifying certain gene groups. Furthermore, it is important to note that these MAGs represent a small fraction of the overall soil microbial diversity which could also bias these findings.

#### Closely-related MAGs both depleted and enriched post-fire, revealing a possible role of glycoside hydrolases

In line with our finding above that success post-fire is not easily predicted by genetic potential or taxonomy, we identified a differential response to fire amongst MAGs belonging to the genus *Aeromicrobium*. Six *Aeromicrobium* MAGs were found to be more relatively abundant in burned vs. unburned (both a week and five months after the burn) while another three were shown to decrease in abundance a week post-burn (**Fig. 5**, purple strip under Actinomycetota). One (T2R4UO.25.metabat2) out of those three decreasing MAGs was found to be enriched in burned vs. unburned five months later. This set of closely related MAGs with contrasting responses to fire offers an interesting opportunity to probe the genetic differences that may explain this differential burn response.

We uncovered that *Aeromicrobium* MAGs enriched in burned vs. unburned a week post-burn had a lower mean predicted doubling time compared to those that were depleted in burned vs. unburned a week post-burn (1.76h vs 3.14h) (see **Supplementary Table 2**). However, we also uncovered intriguing evidence of microbial adaptation to disturbances. Comparative genomics of the MAGs revealed a higher abundance of glycoside hydrolase (GH) enzymes in MAGs depleted a week post-fire (mean of 23) compared to the ones that were enriched (mean of 9) (see **Supplementary Table 2**). GH are enzymes that cleave the glycosidic bonds in complex carbohydrates^86,87^ (e.g., cellulose, starch). It is possible that combustion of polysaccharides might have led to their reduced abundance, as some studies have reported previously^88–90^. In light of this, we hypothesize that MAGs with a higher abundance of GHs are dependent on the degradation of complex carbohydrates, and that a reduction of their availability would impair growth. MAGs with lower abundances of GHs might indicate a more generalist lifestyle in terms of carbohydrate degradation, with less dependence on complex carbohydrates, possibly advantageous in a post-fire condition. Similarly, if the fire releases a pulse of simple carbohydrate-rich dissolved organic carbon, as living plants and microbes are killed and release their cellular contents, these putative copiotrophic MAGs may thrive.

The *Aeromicrobium* MAGs depleted in burned vs. unburned a week post-fire had a much higher abundance of BGCs (mean of 13,144.33 bp vs. 2,216.16 bp when using the proto core region and 53,877.33 bp vs. 5,809.33 bp when using the protocluster region) and a slightly higher abundance of antimicrobial resistance genes (mean of 2.67 vs 0.67) compared to MAGs that were enriched. This would be consistent with reduced competition shortly after a fire (as many microbes die), leading to a selective disadvantage for costly BGCs and their replication, and consistent with our overall finding of MAGs with smaller genome sizes being enriched post-fire. At first glance, it might appear that faster doubling time is the most compelling explanation for differential responses to burning in these nine *Aeromicrobium* MAGs. However, it is important to note that - considering the diversity of microbial interactions in soil, the hypotheses presented here are not mutually exclusive. The fact that one of the MAGs which was depleted in burned vs. unburned (T2R4UO.25.metabat2) a week post-burn but enriched five months later had a comparable number of GHs to MAGs that were enriched at both time points post-burn (see **Supplementary Table 2**), indicates that number of GH enzymes may not be the sole criteria governing the change in microbial abundance post-burn. Our comparative analysis of the GH enzymes opens up new avenues for investigating increases in specific microbial taxa post fire.

## Conclusion

By examining MAGs in detail and, in particular, determining relative coverage of specific MAGs in burned versus unburned timepoint controls, we have uncovered a complex interplay between various traits that increase success post-fire. Although some of these traits were expected based on previous work, our results suggest that many of these are nuanced and perhaps context-dependent. For instance, greater number of spore-related genes was associated with increased post-fire abundance in Bacillota but not in Actinomycetota. Likewise, some genes involved in the biosynthesis of osmotic protectants were actually lower in MAGs increasing post-burn, contrary to expectations. Different taxa often have vastly different metabolic capabilities and requirements, and so it is difficult to compare different species in a metagenome to predict functional differences. However, we did identify a group of MAGs that were closely related but reacted differently to post burn conditions, all belonging to the genus *Aeromicrobium*. In those cases we were able to identify a difference in the number glycosyl hydrolase genes carried, potentially suggesting different degrees of adaptation to using complex carbohydrates as carbon sources. Natural microbiomes are of course difficult to study in a prospective manner, because the majority of members are recalcitrant to culture and therefore cannot be genetically manipulated to directly test the effects of gene knockouts. Therefore, such comparisons between closely-related strains are the next best alternative, but there is still much unknown about how soil microbiomes function and how individual species interact. Although we did unearth several insights into the response of the soil microbiome to fire through genome-resolved methods, the presence and number of genes in specific categories only gives us limited information on bacterial behavior, which is highly dynamic. Given that we found little functional difference in gene profiles on the metagenome level, further insights may require the employment of genome-resolved metatranscriptomics or single-cell methods.

## Methods

### Study region

This study focused on two restored tallgrass prairies in Southwestern Wisconsin located on the University of Wisconsin – Platteville campus (42.726778, −90.494936). Platteville, WI typically has cold winters and warm summers, with mean annual temperatures between 2.1 °C – 13.5 °C and annual precipitation of 917 mm. Rountree prairie is a restored prairie that was previously cultivated. It is located on a relatively flat and low-lying area. Due to this prairie’s low-lying nature, it is historically flooded and has accumulated sediments from upstream topsoils. Restoration efforts began in 1997, and consisted of seeding, invasive species management, and prescribed burns. The vegetation in Rountree prairie is dominated by Canada goldenrod (*Solidago canadensis*), reed canary grass (*Phalaris arundinacea*), and big bluestem (*Andropogon gerardii*). Pioneer prairie is a restored prairie, located on a hillslope with a southern aspect. This prairie has been actively managed since 2003. The vegetation community has a greater species diversity than Rountree prairie, containing predominantly Canada goldenrod (*S. canadensis*), big bluestem (*A. gerardii*), common teasel (*Dipsacus fullonum*), Northern wild senna (*Senna hebecarpa),* and New England aster (*Symphyotrichum novae- angliae*). Both soils are silt loams, had circumneutral pH values (mean 6.6 Pioneer A1 horizon; mean 7.2 Rountree A1 horizon), and moderate C contents (mean 3.8% Pioneer A1 horizon; mean 5.3% Rountree A1 horizon).

### Site assessment methodologies

*Prescribed burn and soil sampling:* In April 2019, team members visited Rountree prairie and Pioneer prairie to begin site assessment, treatment, and sample collection. At each site, six 4 m x 5 m plots were measured, marked, and divided in half into two 2 m x 5 m subplots. A coin flip for each plot determined the side that would receive the burn treatment. Large pieces of wood were sprayed with water and placed on the unburned side (**Fig. 1BC**). Prescribed fire was applied by a backing burn treatment.

Soil samples for each plot were collected after the burned subplot was no longer smoldering – generally within an hour of burning. For each subplot, soil samples were collected and pooled from nine spots, creating a W formation across the plot. The O-horizon, the top 0-5 cm of the A horizon (“A1”) and the 5-15cm of the A horizon (“A2”) were sampled. For each spot, to collect the O horizon, all dead organic material was scooped off the surface of the soil and placed into gallon plastic freezer bags, brushing the surface gently with a gloved hand to collect the organic debris. Within this cleared area, to collect the A1 and A2 horizons, a 2.54 cm diameter soil probe was used to collect 15 cm soil cores, which were split into the top 0-5 cm and the bottom 5-15 cm and combined by depth for each subplot in plastic freezer bags. Samples were stored on ice and moved to a −80 °C freezer within six hours of collection.

Plots were sampled at 7 days, 30 days, and 154 days after the prescribed burn treatment. Soil sampling followed the same procedure as above (but avoiding resampling the exact same locations as previous timepoints). For these subsequent timepoints, sub-samples for DNA extraction were stored in a −20°C freezer until the 154-day timepoint, when all remaining samples were transported to UW-Madison on ice.

### DNA extraction, amplification, and sequencing

O horizon samples were removed from the freezer and ground using a Waring blender prior to sub-sampling for microbial community analyses, in order to collect a sufficiently representative 250 mg sub-sample. DNA extractions were performed for each sample, with one blank extraction every 24 samples (identical methods but using empty tubes), using a DNEasy PowerLyzer PowerSoil DNA extraction kit (QIAGEN, Germantown, MD) following manufacturer’s instructions. About 250 mg of frozen material was weighed for each extraction. Bead beating was performed using a FastPrep-24 5G (MP Biomedicals, LLC, Santa Ana, CA) for 45 seconds at 6 ms^-1^. Extracted DNA was amplified via PCR, with each sample amplified in triplicate, targeting the 16S rRNA gene v4 region (henceforth, “16S”) with 515f and 806r primers^91^. Barcodes and Illumina sequencing adapters were added as per Kozich *et al*.^92^. PCR amplicons were verified for successful amplification and length using gel electrophoresis. The PCR amplicon triplicates were pooled, purified and normalized using a SequalPrep Normalization Plate (96) Kit (ThermoFisher Scientific, Waltham, MA). Samples, including blanks, were pooled and library cleanup was performed using a Wizard SV Gel and PCR Clean- Up System A9282 (Promega, Madison, WI). The pooled library was submitted to the UW Madison Biotechnology Center (UW-Madison, WI) for 2x250 paired end (PE) Illumina MiSeq sequencing for the 16S amplicons. For metagenomic data library preparation and sequencing was done by Novogene with 2x150 PE using Illumina NovaSeq 6000.

### 16S rRNA sequencing analysis

The 16S rRNA gene data was analyzed using Mothur v.1.44.1^93^. Chimeric sequences were identified using the VSEARCH algorithm^94^ and then removed from the dataset. The remaining sequences were classified using the Bayesian classifier against Silva database v138^95^ and those classified as Chloroplast, Mitochondria, unknown, Archaea and Eukaryota were removed. Following this any singletons were removed. Distance matrix (cutoff = 0.04) was created in mothur which was then used to cluster the sequences into Operational Taxonomic Units (OTUs) at distance values of 0.03 and method=opti^96^. Biom file was created using the OTUs clustering at a distance value of 0.03. The Biom file was analyzed using the Phyloseq package^97^ in RStudio.

### Metagenomic analysis

Reads for sample_1 (T2R1BO, 155 Gbp) and sample_2 (T4R1BO, 294 Gbp) were normalized using bbnorm (https://sourceforge.net/projects/bbmap/) with the following parameters, target=70 mindepth=2 prefilter=t. Normalized reads from sample_1 and sample_2 and reads from other 22 samples were assembled using MEGAHIT^98^ with k-list 27,33,43,53,63,73,83,93,103,113,123,133. Metagenomic assembly for some samples was also attempted using metaSPAdes^99^ however, MEGAHIT assemblies were found to be of higher quality, had fewer computational requirements, and was faster. Full full details please refer to **Supplementary Note 2**.

Assembled contigs were binned using Autometa v2^100,101^ (with length cutoff of 3000bp) and MetaBAT2^102^. The large-data-mode of Autometa (max_partition_size=10000) was used for binning samples sample_1 and sample_2. MAGs from Autometa and MetaBAT2 for each sample were combined using DAS Tool^103^ (v1.1.4) with a score threshold of 0.1 to capture MAGs with few single-copy marker genes. MAGs from all the samples were combined and dereplicated using dRep^104^ using options --S_ani 0.98 --length 10000 -comp 50 -con 10. MAGs were dereplicated as 98% ANI to capture as much soil microdiversity as possible. Completeness and contamination metrics generated using checkM2 v1.0.1^51^ were provided using --genomeInfo option to dRep. Taxonomic classification of MAGs was done using GTDB-Tk^105^ (v2.3.2) (database r214). Prokka^106,107^ was used to call and annotate genes in the dereplicated MAGs.

To identify the abundance of each MAGs in each sample all 308 dereplicated bacterial MAGs were combined in a FASTA file and reads from each sample were aligned on it using bowtie2^108^. autometa-coverage entrypoint^101,109,110^ was used to calculate the coverage of each contig. Custom python script was used to calculate the length weighted coverage (ie. abundance) and relative abundance of each MAG in each sample (**Supplementary Table 1C**). To create the heatmap of percentage of bases aligned as shown in **Supplementary Table 1B**, “bed” file generated using bedtools genomecov^110^ during the above coverage analysis by autometa- coverage was used. Number of bases aligned with at least one read were summed and divided by the total lengths of all the contigs in the genome to get the percentage of bases aligned. Bacterial genomes not expected to be present in soil (eg. human-associated or found in extreme environments) were downloaded from NCBI and concatenated in a FASTA file. Reads from each of the 24 soil samples were aligned on the FASTA file and autometa-coverage entrypoint^101,109,110^ was used to calculate the coverage of each contig. Subsequent steps were similar to calculating the percentage of bases aligned on MAGs, where the “bed” file was used for identifying the number of bases aligned with at least one read and the total length of all the contigs.

Contigs (greater than or equal to 3000bp) identified to be bacterial by Autometa were used to identify COG categories using eggNOG-mapper v2.1.9^111^. Coverage of each contig generated by Autometa was used as a proxy for gene coverage.

### Identification of genes involved in fire response traits

Predicted doubling time was calculated using the gRodon2 package^47^ (partial mode) in RStudio using only MAGs with ribosomal proteins equal to or greater than ten as recommended by the author^47^. Prokka annotations were used to identify ribosomal proteins using the grepl command - grepl("ribosomal protein",names(genes),ignore.case = T, perl = TRUE). According to the author of the tool, the doubling time predictions are not accurate for values greater than 5h^47^. To not discard any valuable data, we used 5h as the predicted doubling time in case the doubling time of a MAG was found to be greater than 5h.

Kofamscan^112^, run with -f "detail-tsv", was used to identify the KEGG Ortholog values for the called genes. Annotations with score greater than threshold were chosen as significant hits.In case there were multiple significant hits for a gene, one with the lowest e-value was chosen. In case of identical e-values, the hit with the highest score was chosen. KO values used to identify genes involved in fire response traits can be found in **Supplementary Table 3**.

Spore genes were identified by parsing Distilled and Refined Annotation of Metabolism (DRAM) v1.4.6 (run using the --use_uniref flag)^113,114^ output for spore, sporulation, endospore, sporangia, sporangium and exospore and only taking annotation with ranks A, B or C. Heat shock proteins were identified by parsing DRAM v1.4.6 (using --use_uniref flag)^113,114^ output for heat shock protein and only taking annotation with ranks A, B or C. Glycoside hydrolase genes were identified using the cazy_hits column in DRAM output.

Antimicrobial resistance genes were identified using AMRFinder v3.12.8 (database version 2024-01-31.1) with the --plus flag^115^ and Resistance Gene Identifier v6.0.3 (RGI)^116^ using the Docker image. For AMRFinder following input options were used --nucleotide, --protein, --gff and --annotation_format prokka. For RGI, “main” mode was used and amino acid sequences were provided using Diamond as an alignment_tool. Results for both tools were combined and checked for any duplicates.

For calculating the combined abundance of fire survival traits gene count for HSPs, mycothiol biosynthesis, glutathione biosynthesis, ectoine biosynthesis, glycine betaine biosynthesis, and trehalose biosynthesis were added. Spore genes were added for **Fig. 5F** but not for **Extended Data Fig. 8b**. If annotated the following gene counts were converted to zero before adding as genes for those specific traits are generally not reported in that particular phyla - mycothiol biosynthesis genes in any phyla other than Actinomycetota, and glutathione biosynthesis genes in Actinomycetota and Bacillota.

#### Statistical analysis

Adonis2 function in R (using Bray-Curtis^117^ dissimilarities on relative abundances) was used to conduct the PERMANOVA analysis for PCoA plots.

Significant levels in boxplots comparing the fire response traits in enriched vs. depleted MAGs was calculated using ggsignif R package^118^ using test=“wilcox.test” to conduct the Wilcoxon rank sum test (also known as Mann-Whitney test). Following significance values were used for “map_signif_level” in ggsignif : "***" = 0.001, "**" = 0.01, "*" = 0.05.

## Author contributions

SU: Conceptualization, Data curation, Formal Analysis, Investigation, Methodology, Visualization, Writing – original draft, Writing – review & editing. JW: Data curation, Investigation, Methodology, Writing – review & editing. MY: Conceptualization, Funding acquisition, Investigation, Methodology, Project administration, Resources, Writing – review & editing. CB: Conceptualization, Funding acquisition, Investigation, Methodology, Project administration, Resources, Writing – review & editing. YJ: Conceptualization, Funding acquisition, Investigation, Methodology, Project administration, Resources, Writing – review & editing. AT: Formal Analysis, Resources, Writing – review & editing. CSY: Formal Analysis, Resources, Supervision, Project administration, Writing – review & editing. TW: Conceptualization, Funding acquisition, Investigation, Methodology, Project administration, Resources, Supervision, Writing – review & editing. JCK: Conceptualization, Funding acquisition, Investigation, Methodology, Resources, Supervision, Project administration, Writing – review & editing.

## Data availability

Shotgun metagenomic reads for the 24 samples and the 308 MAGs are uploaded to NCBI under the Bioproject PRJNA1109856.

## Acknowledgments

We want to thank Penguin computing (https://www.penguinsolutions.com/) for letting us use their hardware for metagenome assembly analysis. The authors thank Dr. Timothy Berry, a team of UW-Platteville undergraduate students from Soil and Crop Science program (Mr. Garrett Larsen, Ms. Jessica Helwig, Mr. Joseph Creanza, Ms. Kaylee Finseth, Ms. Courtney Unzicker, Ms. Hannah McWhirter, Mr. Austin Benzing, Mr. Sara Raemisch, Ms. Caroline Sawyer, Mr. Charles Peterson, Ms. Katie Walsh), and student volunteers from Ecological Restoration and Resource Management program for their assistance with conducting burns and soil sampling. The authors also wish to thank Karthik Anantharaman for valuable insights during the preparation of this paper. The field experiment was funded by a grant to TW, MV, YBJ, and CB from the University of Wisconsin Consortium for Extension and Research in Agriculture and Natural Resources. TW was supported by an NSF CAREER grant during data analysis and paper writing (NSF-2045864).

---

**Extended Data Fig. 1.** Shannon diversity of OTU relative abundance for Pioneer prairie at a) A1 horizon and b) A2 horizon. Shannon diversity of OTU relative abundance for Rountree prairie at c) A1 horizon and d) A2 horizon.

**Extended Data Fig. 2.** PCoA ordination of Bray Curtis dissimilarities for OTU relative abundance for Pioneer prairie at a) A1 horizon and b) A2 horizon. PCoA ordination of Bray Curtis dissimilarities for OTU relative abundance for Rountree prairie at c) A1 horizon and d) A2 horizon.

**Extended Data Fig. 3.** PCoA (using Bray-Curtis dissimilarities) of relative abundance of COG counts for a) a week after the burn and b) five months after the burn. PCoA (using Bray-Curtis dissimilarities) of relative abundance of COGs coverage c) a week after the burn and d) five months after the burn in the O horizon at the Roundtree Prairie

**Extended Data Fig. 4.** PCoA (using Bray-Curtis dissimilarities) of relative abundance of MAGs, a) A week after burn and b) Five months after the burn in the Rountree Prairie.

**Extended Data Fig. 5.** Characteristics for all MAGs classified by whether they are depleted or enriched in burned vs. unburned plots one week and five months post-fire from rountree prairie. a) Genome size of MAGs with greater than 70% completeness. b) Length weighted GC%. c) Spore gene count, excluding Bacillota MAGs. d) Spore gene count, only Actinomycetota MAGs. NS indicates no significant difference between the groups. * indicates significance at p < 0.05, ** indicates significance at p < 0.01, and *** indicates significance at p < 0.001.

**Extended Data Fig. 6.** Benzoate degradation pathway. Genes highlighted in green were present in all four MAGs found to increase after the burn. *pcaIJ* was present in MAGs T2R1BO.metabat2.154 and T4R3BO.57.metabat2 (order *Burkholderiales,* family *Burkholderiaceae_B* and *Burkholderiaceae* respectively), *pcaF* was present in MAGs T2R5BO_autometa_bin_0008 and TT2R6UO_autometa_bin_0026 (genus *Pseudomonas_E*, family *Pseudomonadaceae*) and *fadA* was present only in T4R3BO.57.metabat2 (*Duganella* sp.). Pathway adapted from KEGG database.

**Extended Data Fig. 7.** Characteristics for all MAGs classified by whether they are depleted or enriched in burned vs. unburned plots one week and five months post-fire from rountree prairie. Number of genes for a) heat shock proteins, b) glutathione biosynthesis, excluding Bacillota and Actinomycetota MAGs, c) ectoine biosynthesis and d) glycine betaine biosynthesis. NS indicates no significant difference between the groups. * indicates significance at p < 0.05, ** indicates significance at p < 0.01, and *** indicates significance at p < 0.001.

**Extended Data Fig. 8.** Gene count for all fire survival traits combined. MAGs are classified by whether they are depleted or enriched in burned vs. unburned plots one week and five months post-fire from rountree prairie. Scale on y-axis is reduced by a factor of 5 after 60. NS indicates no significant difference between the groups. * indicates significance at p < 0.05, ** indicates significance at p<0.01, and *** indicates significance at p < 0.001.

## References

1. Schnitzer, S. A. et al. Soil microbes drive the classic plant diversity-productivity pattern. Ecology 92, 296–303 (2011).

2. van der Heijden, M. G. A., Bardgett, R. D. & van Straalen, N. M. The unseen majority: soil microbes as drivers of plant diversity and productivity in terrestrial ecosystems. Ecol. Lett. 11, 296–310 (2008).

3. Crowther, T. W. et al. The global soil community and its influence on biogeochemistry. Science 365, eaav0550 (2019).

4. Ramirez, K. S. et al. Biogeographic patterns in below-ground diversity in New York City’s Central Park are similar to those observed globally. Proc. R. Soc. B. 281, 20141988 (2014).

5. Bardgett, R. D. & van der Putten, W. H. Belowground biodiversity and ecosystem functioning. Nature 515, 505–511 (2014).

6. Howe, A. C. et al. Tackling soil diversity with the assembly of large, complex metagenomes. Proc. Natl. Acad. Sci. U. S. A. 111, 4904–4909 (2014).

7. Ryan, K. C., Knapp, E. E. & Varner, J. M. Prescribed fire in North American forests and woodlands: history, current practice, and challenges. Front. Ecol. Environ. 11, e15–e24 (2013).

8. Wu, X., Sverdrup, E., Mastrandrea, M. D., Wara, M. W. & Wager, S. Low-intensity fires mitigate the risk of high-intensity wildfires in California’s forests. Sci Adv 9, eadi4123 (2023).

9. Hmielowski, T. L. et al. Prioritizing land management efforts at a landscape scale: a case study using prescribed fire in Wisconsin. Ecol. Appl. 26, 1018–1029 (2016).

10. Nelson, A. R. et al. Wildfire-dependent changes in soil microbiome diversity and function. Nat Microbiol 7, 1419–1430 (2022).

11. Holden, S. R. & Treseder, K. K. A meta-analysis of soil microbial biomass responses to forest disturbances. Front. Microbiol. 4, 163 (2013).

12. Dooley, S. R. & Treseder, K. K. The effect of fire on microbial biomass: a meta-analysis of field studies. Biogeochemistry 109, 49–61 (2012).

13. Barreiro, A. & Díaz-Raviña, M. Fire impacts on soil microorganisms: mass, activity, and diversity. Curr. Opin. Environ. Sci. Health 22, 100264 (2021).

14. Sáenz de Miera L. E., Pinto, R., Gutierrez-Gonzalez, J. J., Calvo, L. & Ansola, G. Wildfire effects on diversity and composition in soil bacterial communities. Sci. Total Environ. 726, 138636 (2020).

15. Whitman, T. et al. Soil bacterial and fungal response to wildfires in the Canadian boreal forest across a burn severity gradient. Soil Biol. Biochem. 138, 107571 (2019).

16. Fernández-González, A. J. et al. The rhizosphere microbiome of burned holm-oak: potential role of the genus *Arthrobacter* in the recovery of burned soils. Sci. Rep. 7, 6008 (2017).

17. Soria, R. et al. Short-term response of soil bacterial communities after prescribed fires in semi-arid Mediterranean forests. Fire 6, 145 (2023).

18. Villadas, P. J. et al. The soil microbiome of the laurel forest in Garajonay National Park (La Gomera, Canary Islands): comparing unburned and burned habitats after a wildfire. Forests 10, 1051 (2019).

19. Chen, H. et al. Functional redundancy in soil microbial community based on metagenomics across the globe. Front. Microbiol. 13, 878978 (2022).

20. Louca, S. et al. Function and functional redundancy in microbial systems. Nat Ecol Evol 2, 936–943 (2018).

21. Johnson, D. B., Woolet, J., Yedinak, K. M. & Whitman, T. Experimentally determined traits shape bacterial community composition one and five years following wildfire. Nat Ecol Evol 7, 1419–1431 (2023).

22. Nemergut, D. R. et al. Decreases in average bacterial community rRNA operon copy number during succession. ISME J. 10, 1147–1156 (2016).

23. Adkins, J., Docherty, K. M. & Miesel, J. R. Copiotrophic bacterial traits increase with burn severity one year after a wildfire. Front. For. Glob. Chang. 5, 873527 (2022).

24. Bickel, S. & Or, D. The chosen few-variations in common and rare soil bacteria across biomes. ISME J. 15, 3315–3325 (2021).

25. Bowers, R. M. et al. Minimum information about a single amplified genome (MISAG) and a metagenome-assembled genome (MIMAG) of bacteria and archaea. Nat Biotechnol 35, 725–731 (2017).

26. Fernández-González, A. J. et al. Long-term persistence of three microbial wildfire biomarkers in forest soils. Forests 14, 1383 (2023).

27. Weber, C. F., Lockhart, J. S., Charaska, E., Aho, K. & Lohse, K. A. Bacterial composition of soils in ponderosa pine and mixed conifer forests exposed to different wildfire burn severity. Soil Biol. Biochem. 69, 242–250 (2014).

28. Bowman, J. P. Genome-wide and constrained ordination-based analyses of EC code data support reclassification of the species of Massilia La Scola et al. 2000 into Telluria Bowman et al. 1993, Mokoshia gen. nov. and Zemynaea gen. nov. Int. J. Syst. Evol. Microbiol. 73, 005991 (2023).

29. Parks, D. H. et al. A standardized bacterial taxonomy based on genome phylogeny substantially revises the tree of life. Nat. Biotechnol. 36, 996–1004 (2018).

30. Brown, C. T. et al. Unusual biology across a group comprising more than 15% of domain Bacteria. Nature 523, 208–211 (2015).

31. Castelle, C. J. et al. Biosynthetic capacity, metabolic variety and unusual biology in the CPR and DPANN radiations. Nat. Rev. Microbiol. 16, 629–645 (2018).

32. He, X. et al. Cultivation of a human-associated TM7 phylotype reveals a reduced genome and epibiotic parasitic lifestyle. Proc. Natl. Acad. Sci. U. S. A. 112, 244–249 (2015).

33. Moreira, D., Zivanovic, Y., López-Archilla, A. I., Iniesto, M. & López-García, P. Reductive evolution and unique predatory mode in the CPR bacterium *Vampirococcus lugosii*. Nat. Commun. 12, 2454 (2021).

34. Yakimov, M. M. et al. Cultivation of a vampire: ‘*Candidatus* Absconditicoccus praedator’. Environ. Microbiol. 24, 30–49 (2022).

35. Batinovic, S., Rose, J. J. A., Ratcliffe, J., Seviour, R. J. & Petrovski, S. Cocultivation of an ultrasmall environmental parasitic bacterium with lytic ability against bacteria associated with wastewater foams. Nat Microbiol 6, 703–711 (2021).

36. Gong, J., Qing, Y., Guo, X. & Warren, A. ‘*Candidatus* Sonnebornia yantaiensis’, a member of candidate division OD1, as intracellular bacteria of the ciliated protist *Paramecium bursaria* (Ciliophora, Oligohymenophorea). Syst. Appl. Microbiol. 37, 35–41 (2014).

37. Kuroda, K. et al. Symbiosis between *Candidatus* Patescibacteria and archaea discovered in wastewater-treating bioreactors. MBio 13, e01711–22 (2022).

38. Kuroda, K. et al. Novel cross-domain symbiosis between *Candidatus* Patescibacteria and hydrogenotrophic methanogenic archaea *Methanospirillum* discovered in a methanogenic ecosystem. Microbes Environ. 37, ME22063 (2022).

39. Kuroda, K. et al. Microscopic and metatranscriptomic analyses revealed unique cross-domain parasitism between phylum *Candidatus* Patescibacteria/candidate phyla radiation and methanogenic archaea in anaerobic ecosystems. MBio 15, e03102–23 (2024).

40. Chen, X. et al. ‘*Candidatus* Nealsonbacteria’ are likely biomass recycling ectosymbionts of methanogenic archaea in a stable benzene-degrading enrichment culture. Appl. Environ. Microbiol. 89, e00025–23 (2023).

41. Fischer, M. S., Patel, N. J., de Lorimier, P. J. & Traxler, M. F. Prescribed fire selects for a pyrophilous soil sub-community in a northern California mixed conifer forest. Environ. Microbiol. 25, 2498–2515 (2023).

42. Zeba, N., Berry, T. D., Fischer, M. S., Traxler, M. F. & Whitman, T. Soil carbon mineralization and microbial community dynamics in response to pyrogenic organic matter addition. Soil Biol. Biochem. 191, 109328 (2024).

43. Blin, K. et al. antiSMASH 7.0: new and improved predictions for detection, regulation, chemical structures and visualisation. Nucleic Acids Res. 51, W46–W50 (2023).

44. Crits-Christoph, A., Diamond, S., Butterfield, C. N., Thomas, B. C. & Banfield, J. F. Novel soil bacteria possess diverse genes for secondary metabolite biosynthesis. Nature 558, 440–444 (2018).

45. Charlop-Powers, Z., Owen, J. G., Reddy, B. V. B., Ternei, M. A. & Brady, S. F. Chemical- biogeographic survey of secondary metabolism in soil. Proc. Natl. Acad. Sci. U. S. A. 111, 3757–3762 (2014).

46. Ma, B. et al. A genomic catalogue of soil microbiomes boosts mining of biodiversity and genetic resources. Nat. Commun. 14, 7318 (2023).

47. Weissman, J. L., Hou, S. & Fuhrman, J. A. Estimating maximal microbial growth rates from cultures, metagenomes, and single cells via codon usage patterns. Proc. Natl. Acad. Sci. U. S. A. 118, e2016810118 (2021).

48. Piton, G. et al. Life history strategies of soil bacterial communities across global terrestrial biomes. Nat Microbiol 8, 2093–2102 (2023).

49. Liu, H., et al. Warmer and drier ecosystems select for smaller bacterial genomes in global soils. Imeta 2, e70 (2023).

50. Simonsen, A. K. Environmental stress leads to genome streamlining in a widely distributed species of soil bacteria. ISME J. 16, 423–434 (2022).

51. Chklovski, A., Parks, D. H., Woodcroft, B. J. & Tyson, G. W. CheckM2: a rapid, scalable and accurate tool for assessing microbial genome quality using machine learning. Nat. Methods 20, 1203–1212 (2023).

52. Yakovchuk, P., Protozanova, E. & Frank-Kamenetskii, M. D. Base-stacking and base- pairing contributions into thermal stability of the DNA double helix. Nucleic Acids Res. 34, 564–574 (2006).

53. Reisser, M., Purves, R. S., Schmidt, M. W. I. & Abiven, S. Pyrogenic carbon in soils: a literature-based inventory and a global estimation of its content in soil organic carbon and stocks. Front. Earth Sci. 4, 80 (2016).

54. Keiluweit, M., Nico, P. S., Johnson, M. G. & Kleber, M. Dynamic molecular structure of plant biomass-derived black carbon (biochar). Environ. Sci. Technol. 44, 1247–1253 (2010).

55. Zhang, L. et al. Habitat heterogeneity induced by pyrogenic organic matter in wildfire- perturbed soils mediates bacterial community assembly processes. ISME J. 15, 1943–1955 (2021).

56. Fuchs, G., Boll, M. & Heider, J. Microbial degradation of aromatic compounds - from one strategy to four. Nat. Rev. Microbiol. 9, 803–816 (2011).

57. Nicholson, W. L., Munakata, N., Horneck, G., Melosh, H. J. & Setlow, P. Resistance of *Bacillus* endospores to extreme terrestrial and extraterrestrial environments. Microbiol. Mol. Biol. Rev. 64, 548–572 (2000).

58. Setlow, P. Spore resistance properties. Microbiol Spectr **2**, TBS–0003–2012 (2014).

59. Muñoz-Dorado, J., Marcos-Torres, F. J., García-Bravo, E., Moraleda-Muñoz, A. & Pérez, J. Myxobacteria: moving, killing, feeding, and surviving together. Front. Microbiol. 7, 781 (2016).

60. Kaplan-Levy, R. N., Hadas, O., Summers, M. L., Rücker, J. & Sukenik, A. Akinetes: dormant cells of cyanobacteria. in Dormancy and Resistance in Harsh Environments (eds. Lubzens, E., Cerda, J. & Clark, M.) vol. 12 5–27 (Springer Berlin Heidelberg, 2010).

61. Yabe, S., Aiba, Y., Sakai, Y., Hazaka, M. & Yokota, A. A life cycle of branched aerial mycelium- and multiple budding spore-forming bacterium *Thermosporothrix hazakensis* belonging to the phylum Chloroflexi. J. Gen. Appl. Microbiol. 56, 137–141 (2010).

62. Beskrovnaya, P., Sexton, D. L., Golmohammadzadeh, M., Hashimi, A. & Tocheva, E. I. Structural, metabolic and evolutionary comparison of bacterial endospore and exospore formation. Front. Microbiol. 12, 630573 (2021).

63. McKenney, P. T., Driks, A. & Eichenberger, P. The Bacillus subtilis endospore: assembly and functions of the multilayered coat. Nat. Rev. Microbiol. 11, 33–44 (2013).

64. McCormick, J. R. & Flärdh, K. Signals and regulators that govern Streptomyces development. FEMS Microbiol. Rev. 36, 206–231 (2012).

65. Beskrovnaya, P. et al. No endospore formation confirmed in members of the phylum Proteobacteria. Appl. Environ. Microbiol. 87, e02312–20 (2021).

66. Traag, B. A. et al. Do mycobacteria produce endospores? Proc. Natl. Acad. Sci. U. S. A. 107, 878–881 (2010).

67. Pérez-Valera, E., Goberna, M. & Verdú, M. Fire modulates ecosystem functioning through the phylogenetic structure of soil bacterial communities. Soil Biol. Biochem. 129, 80–89 (2019).

68. Lucas-Borja, M. E. et al. Immediate fire-induced changes in soil microbial community composition in an outdoor experimental controlled system. Sci. Total Environ. 696, 134033 (2019).

69. Prendergast-Miller, M. T. et al. Wildfire impact: natural experiment reveals differential short- term changes in soil microbial communities. Soil Biol. Biochem. 109, 1–13 (2017).

70. Hinojosa, M. B., Laudicina, V. A., Parra, A., Albert-Belda, E. & Moreno, J. M. Drought and its legacy modulate the post-fire recovery of soil functionality and microbial community structure in a Mediterranean shrubland. Glob. Chang. Biol. 25, 1409–1427 (2019).

71. Cobo-Díaz, J. F. et al. Metagenomic assessment of the potential microbial nitrogen pathways in the rhizosphere of a mediterranean forest after a wildfire. Microb. Ecol. 69, 895–904 (2015).

72. Agbeshie, A. A., Abugre, S., Atta-Darkwa, T. & Awuah, R. A review of the effects of forest fire on soil properties. *J*. For. Res. 33, 1419–1441 (2022).

73. Dove, N. C., Taş, N. & Hart, S. C. Ecological and genomic responses of soil microbiomes to high-severity wildfire: linking community assembly to functional potential. ISME J. 16, 1853–1863 (2022).

74. Sigmund, G. et al. Environmentally persistent free radicals are ubiquitous in wildfire charcoals and remain stable for years. Commun Earth Environ 2, 68 (2021).

75. Abraham, J., Dowling, K. & Florentine, S. The unquantified risk of post-fire metal concentration in soil: a review. Water Air Soil Pollut. 228, 175 (2017).

76. Stankov Jovanovic, V. P., et al. Wild fire impact on copper, zinc, lead and cadmium distribution in soil and relation with abundance in selected plants of Lamiaceae family from Vidlic Mountain (Serbia). Chemosphere 84, 1584–1591 (2011).

77. Ulrich, K. & Jakob, U. The role of thiols in antioxidant systems. Free Radic. Biol. Med. 140, 14–27 (2019).

78. Fahey, R. C., Brown, W. C., Adams, W. B. & Worsham, M. B. Occurrence of glutathione in bacteria. J. Bacteriol. 133, 1126–1129 (1978).

79. Newton, G. L. et al. Bacillithiol is an antioxidant thiol produced in Bacilli. Nat. Chem. Biol. 5, 625–627 (2009).

80. Newton, G. L., Buchmeier, N. & Fahey, R. C. Biosynthesis and functions of mycothiol, the unique protective thiol of Actinobacteria. Microbiol. Mol. Biol. Rev. 72, 471–494 (2008).

81. Santín, C. & Doerr, S. H. Fire effects on soils: the human dimension. Phil. Trans. R. Soc. B 371, 20150171 (2016).

82. Reina-Bueno, M. et al. Role of trehalose in heat and desiccation tolerance in the soil bacterium *Rhizobium etli*. BMC Microbiol. 12, 207 (2012).

83. Wood, J. M. Bacterial osmoregulation: a paradigm for the study of cellular homeostasis. Annu. Rev. Microbiol. 65, 215–238 (2011).

84. Czech, L. et al. Role of the extremolytes ectoine and hydroxyectoine as stress protectants and nutrients: genetics, phylogenomics, biochemistry, and structural analysis. Genes 9, 177 (2018).

85. Iturriaga, G., Suárez, R. & Nova-Franco, B. Trehalose metabolism: from osmoprotection to signaling. Int. J. Mol. Sci. 10, 3793–3810 (2009).

86. Lombard, V., Golaconda Ramulu, H., Drula, E., Coutinho, P. M. & Henrissat, B. The carbohydrate-active enzymes database (CAZy) in 2013. Nucleic Acids Res. 42, D490– D495 (2014).

87. Henrissat, B. & Davies, G. Structural and sequence-based classification of glycoside hydrolases. Curr. Opin. Struct. Biol. 7, 637–644 (1997).

88. Dymov, A. A. et al. Soils and soil organic matter transformations during the two years after a low-intensity surface fire (Subpolar Ural, Russia). Geoderma 404, 115278 (2021).

89. Miltner, A. & Zech, W. Effects of minerals on the transformation of organic matter during simulated fire-induced pyrolysis. Org. Geochem. 26, 175–182 (1997).

90. Martín, A., Díaz-Raviña, M. & Carballas, T. Evolution of composition and content of soil carbohydrates following forest wildfires. Biol. Fertil. Soils 45, 511–520 (2009).

91. Walters, W. et al. Improved bacterial 16S rRNA gene (V4 and V4-5) and fungal internal transcribed spacer marker gene primers for microbial community surveys. mSystems 1, e00009–15 (2016).

92. Kozich, J. J., Westcott, S. L., Baxter, N. T., Highlander, S. K. & Schloss, P. D. Development of a dual-index sequencing strategy and curation pipeline for analyzing amplicon sequence data on the MiSeq Illumina sequencing platform. Appl. Environ. Microbiol. 79, 5112–5120 (2013).

93. Schloss, P. D. et al. Introducing mothur: open-source, platform-independent, community-supported software for describing and comparing microbial communities. Appl. Environ. Microbiol. 75, 7537–7541 (2009).

94. Rognes, T., Flouri, T., Nichols, B., Quince, C. & Mahé, F. VSEARCH: a versatile open source tool for metagenomics. PeerJ 4, e2584 (2016).

95. Quast, C. et al. The SILVA ribosomal RNA gene database project: improved data processing and web-based tools. Nucleic Acids Res. 41, D590–D596 (2013).

96. Westcott, S. L. & Schloss, P. D. OptiClust, an improved method for assigning amplicon-based sequence data to operational taxonomic units. mSphere 2, e00073–17 (2017).

97. McMurdie, P. J. & Holmes, S. phyloseq: an R package for reproducible interactive analysis and graphics of microbiome census data. PLoS One 8, e61217 (2013).

98. Li, D., Liu, C.-M., Luo, R., Sadakane, K. & Lam, T.-W. MEGAHIT: an ultra-fast single-node solution for large and complex metagenomics assembly via succinct *de Bruijn* graph. Bioinformatics 31, 1674–1676 (2015).

99. Nurk, S., Meleshko, D., Korobeynikov, A. & Pevzner, P. A. metaSPAdes: a new versatile metagenomic assembler. Genome Res. 27, 824–834 (2017).

100. Miller, I. J. et al. Autometa: automated extraction of microbial genomes from individual shotgun metagenomes. Nucleic Acids Res. 47, e57 (2019).

101. Rees, E. R. et al. Autometa 2: A versatile tool for recovering genomes from highly-complex metagenomic communities. doi:Preprint at https://www.biorxiv.org/content/10.1101/2023.09.01.555939v1 (2023).

102. Kang, D. D. et al. MetaBAT 2: an adaptive binning algorithm for robust and efficient genome reconstruction from metagenome assemblies. PeerJ 7, e7359 (2019).

103. Sieber, C. M. K. et al. Recovery of genomes from metagenomes via a dereplication, aggregation and scoring strategy. Nat Microbiol 3, 836–843 (2018).

104. Olm, M. R., Brown, C. T., Brooks, B. & Banfield, J. F. dRep: a tool for fast and accurate genomic comparisons that enables improved genome recovery from metagenomes through de-replication. ISME J. 11, 2864–2868 (2017).

105. Chaumeil, P.-A., Mussig, A. J., Hugenholtz, P. & Parks, D. H. GTDB-Tk v2: memory friendly classification with the genome taxonomy database. Bioinformatics 38, 5315–5316 (2022).

106. Seemann, T. Prokka: rapid prokaryotic genome annotation. Bioinformatics 30, 2068–2069 (2014).

107. Hyatt, D. et al. Prodigal: prokaryotic gene recognition and translation initiation site identification. BMC Bioinformatics 11, 119 (2010).

108. Langmead, B. & Salzberg, S. L. Fast gapped-read alignment with Bowtie 2. Nat. Methods 9, 357–359 (2012).

109. Li, H. et al. The sequence alignment/map format and SAMtools. Bioinformatics 25, 2078– 2079 (2009).

110. Quinlan, A. R. & Hall, I. M. BEDTools: a flexible suite of utilities for comparing genomic features. Bioinformatics 26, 841–842 (2010).

111. Cantalapiedra, C. P., Hernández-Plaza, A., Letunic, I., Bork, P. & Huerta-Cepas, J. eggNOG-mapper v2: Functional annotation, orthology assignments, and domain prediction at the metagenomic scale. Mol. Biol. Evol. 38, 5825–5829 (2021).

112. Aramaki, T. et al. KofamKOALA: KEGG Ortholog assignment based on profile HMM and adaptive score threshold. Bioinformatics 36, 2251–2252 (2020).

113. Shaffer, M. et al. DRAM for distilling microbial metabolism to automate the curation of microbiome function. Nucleic Acids Res. 48, 8883–8900 (2020).

114. Wang, Y. et al. A crowdsourcing open platform for literature curation in UniProt. PLoS Biol. 19, e3001464 (2021).

115. Feldgarden, M. et al. AMRFinderPlus and the reference gene catalog facilitate examination of the genomic links among antimicrobial resistance, stress response, and virulence. Sci. Rep. 11, 12728 (2021).

116. Alcock, B. P. et al. CARD 2023: expanded curation, support for machine learning, and resistome prediction at the Comprehensive Antibiotic Resistance Database. Nucleic Acids Res. 51, D690–D699 (2023).

117. Bray, J. R. & Curtis, J. T. An ordination of the upland forest communities of southern Wisconsin. Ecol. Monogr. 27, 325–349 (1957).

118. Ahlmann-Eltze, C. & Patil, I. ggsignif: R package for displaying significance brackets for ‘ggplot2’. doi:Preprint at https://osf.io/preprints/psyarxiv/7awm6 (2021).

