## Supplementary Note 1 for "A granular view of the traits controlling soil bacterial community reconstruction following a prescribed burn"

### Supplementary Note 1. Sequencing depth estimation for shotgun metagenomics

Given the high microbial diversity of soil, we aimed to sequence samples with sufficient depth to capture as many novel genomes as possible. We hypothesized that greater coverage would capture more of the soil biodiversity and yield a higher number of high-quality metagenome-assembled genomes (MAGs).

Initially, two samples were subjected to deep shotgun sequencing. To scale these sequencing runs, 12 publicly available soil metagenomes were examined across a wide range of sequencing depths (**Table SI 1**) using Nonpareil 3[1] (kmer mode). Based on recent work by Rodriguez-R *et al.*[1], who demonstrated a relationship between 16S Shannon diversity and Nonpareil-measured diversity in shotgun datasets, 90% metagenomic coverage was targeted for sample_1 and sample_2, with study design IDs of T2R1BO and T4R1BO respectively (**Supplementary Table 1A**), by sequencing them at depths of 155 Gbp and 294 Gbp, respectively. However, upon sequencing, Nonpareil diversity was higher than expected (23.6 and 24.8), leading to coverages of 69.29% and 63.69%. Assembly of these datasets also did not yield as many high-quality MAGs as expected, only 5 for each sample_1 and sample_2. Therefore, the strategy was changed to sequence more samples at a lower depth rather than a few at very high depth, as it was hypothesized that sampling closely in time and space would yield different high abundance MAGs in different samples which would nevertheless be conserved in the broader sample set. Therefore the remaining burned and unburned samples of Rountree prairie a week and five months post-burn (22 samples) were sequenced at lower depth than the initial two samples.

**Table SI 1.** Sequencing coverage estimates for metagenomes using Nonpareil 3. SeqKit[2] was used to calculate the number of bases in a metagenome.

| Sample | Sequencing Depth | Coverage (%) | Nonpareil diversity |
|---|---|---|---|
| Sample_1 (T2R1BO) | 155.2G bases | 69.29 | 23.6 |
| Sample_2 (T4R1BO) | 294.3G bases | 63.69 | 24.8 |
| SRR7042366 | 9.6G bases | 19.93 | 23.93 |
| SRR7042342 | 21G bases | 37.98 | 23.6 |
| SRR7042381 | 84.3G bases | 52.27 | 23.95 |
| SRR8439256 | 100G bases | 89.42 | 21.53 |
| ERR3697022 | 115.9G bases | 63.32 | 23.64 |
| SRR8439259 | 128.1G bases, | 90.75 | 21.51 |
| SRR9009774 | 153.1G bases | 61.66 | 24.07 |
| SRR9009773 | 157.5G bases | 57.81 | 24.45 |
| SRR8554900 | 163.5G bases | 71.84 | 23.51 |
| SRR9007738 | 193.9G bases | 91.40 | 22.69 |
| SRR10389008 | 201.1G bases | 93.12 | 22.55 |
| SRR15170210 | 356.5G bases | 86.68 | 22.79 |

### Supplementary Note 2. Comparison between MEGAHIT and metaSPADes for soil metagenome assembly

Based on our experience with assembling host-associated metagenomes, we have repeatedly found metaSPAdes[3] assembly statistics to be of higher quality (longer contigs, better N50, etc) compared to MEGAHIT[4,5]. However, these better assembly statistics come at the cost of longer runtime and higher RAM requirements. Our initial attempt to assemble our soil datasets using metaSPAdes was unsuccessful, as it required more RAM than the 1 TB we had available. To compare the assembly quality between metaSPAdes[3] and MEGAHIT[4,5], we collaborated with MemVerge Inc (https://memverge.com/). MemVerge's software uses persistent memory (PMem) as an expansion to dynamic RAM (DRAM) (the type of RAM used in modern computers). It enables the use of both DRAM and PMem to run memory-expensive jobs quicker and cheaper than using DRAM alone. By combining Memverge's Memory Machine 2.5.2 with Penguin Computing's (https://www.penguinsolutions.com/) hardware (5220 CPUs, 786 GB DRAM, and 3 TB PMEM), we were able to assemble three soil datasets referred to as sample_3 (63.7 Gbp), sample_4 (48.7 Gbp) and sample_5, (60.5 Gbp) with study design IDs of T2R1UO, T2R2BO and T4R1UO respectively

Despite the longer runtime, metaSPAdes had poor assembly quality - smaller number of long (>1kbp) and super long contigs (>50 kbp) - for all three of the test soil datasets as compared to MEGAHIT **(Table SI 2)**. Furthermore, while the successful assembly of the datasets using metaSPAdes required a combination of PMem and DRAM, MEGAHIT was able to finish the assemblies with just DRAM (< 1 TB). Even though metaSPAdes had a much longer assembly, most of the contigs were below 1000 bp, and hence are usually filtered out by the majority of the available binning algorithms. Assembly of the larger sample_1 (155 Gbp) and sample_2 (294 Gbp) datasets discussed above were attempted; however, even with enough RAM, initial estimates predicted the process would take more than a month. As an example, just the read correction step for the sample_2 dataset took about 18 days (using 72 threads with hyperthreading) and just the K21 assembly step for sample_1 was unfinished after nine days (using 36 threads). Given these practical considerations, the remaining metagenomes were assembled with MEGAHIT.

MetaQUAST[6,7] (v5.0.2) with flags --min-contig 1 --max-ref-num 1 was used to measure the assembly statistics mentioned in **Table SI 2**. Since only statistics without reference were reported the number of references were kept at 1 to minimize runtime and conserve disk space.

**Table SI 2.** Sequencing coverage estimates for metagenomes using Nonpareil 3. metaSPADes was run using Memverge's Memory Machine on Penguin Computing's hardware, whereas MEGAHIT was run on our own local server. *Using hyperthreading.

| **Statistic** | **Sample_3 (63.7 Gbp, T2R1UO)** | | **Sample_4 (48.7 Gbp, T2R2BO)** | | **Sample_5 (60.5 Gbp, T4R1UO)** | |
|---|---|---|---|---|---|---|
| | **MEGAHIT** | **metaSPADes** | **MEGAHIT** | **metaSPADes** | **MEGAHIT** | **metaSPADes** |
| Runtime (h) | 44.71 | 228.53 | 34 | 113.99 | 21.48 | 175.74 |
| Threads used | 25 | 72* | 25 | 36 | 30 | 72* |
| Total contigs (>=0 bp) | 11,043,691 | 33,506,904 | 7,271,736 | 22,850,760 | 8,403,482 | 35,067,587 |

| Statistic | Sample_3 (63.7 Gbp, T2R1UO) | | Sample_4 (48.7 Gbp, T2R2BO) | | Sample_5 (60.5 Gbp, T4R1UO) | |
|---|---|---|---|---|---|---|
| | MEGAHIT | metaSPADes | MEGAHIT | metaSPADes | MEGAHIT | metaSPADes |
| Contigs >=1000 bp | 1,041,505 | 588,962 | 703,865 | 422,928 | 558,846 | 302,534 |
| Contigs >=10000 bp | 3,878 | 3,033 | 3,530 | 2,415 | 1,244 | 1,086 |
| Contigs >=50000 bp | 18 | 11 | 30 | 15 | 18 | 19 |
| Total length (>= 0 bp) | 6,720,469,839 | 10,892,333,786 | 4,396,948,817 | 7,393,285,757 | 4,711,397,753 | 10,722,045,557 |
| Length >=1000 bp | 1,783,953,103 | 1,040,457,845 | 1,245,909,669 | 772,294,772 | 863,684,215 | 493,240,769 |
| Length >=10000 bp | 57,723,221 | 46,095,510 | 54,682,596 | 38,013,481 | 20,885,036 | 19,008,753 |

| Statistic | Sample_3 (63.7 Gbp, T2R1UO) | | Sample_4 (48.7 Gbp, T2R2BO) | | Sample_5 (60.5 Gbp, T4R1UO) | |
|---|---|---|---|---|---|---|
| | **MEGAHIT** | **metaSPADes** | **MEGAHIT** | **metaSPADes** | **MEGAHIT** | **metaSPADes** |
| Length >=50000 bp | 1,158,215 | 669,174 | 1,959,369 | 1,175,159 | 1,227,812 | 1,273,525 |

### References

1. Rodriguez-R, L. M., Gunturu, S., Tiedje, J. M., Cole, J. R. & Konstantinidis, K. T. Nonpareil 3: fast estimation of metagenomic coverage and sequence diversity. *mSystems* **3**, e00039–18 (2018).
2. Shen, W., Le, S., Li, Y. & Hu, F. SeqKit: a cross-platform and ultrafast toolkit for FASTA/Q file manipulation. *PLoS One* **11**, e0163962 (2016).
3. Nurk, S., Meleshko, D., Korobeynikov, A. & Pevzner, P. A. metaSPAdes: a new versatile metagenomic assembler. *Genome Res.* **27**, 824–834 (2017).
4. Li, D. *et al.* MEGAHIT v1.0: A fast and scalable metagenome assembler driven by advanced methodologies and community practices. *Methods* **102**, 3–11 (2016).
5. Li, D., Liu, C.-M., Luo, R., Sadakane, K. & Lam, T.-W. MEGAHIT: an ultra-fast single-node solution for large and complex metagenomics assembly via succinct *de Bruijn* graph. *Bioinformatics* **31**, 1674–1676 (2015).
6. Mikheenko, A., Saveliev, V. & Gurevich, A. MetaQUAST: evaluation of metagenome assemblies. *Bioinformatics* **32**, 1088–1090 (2016).
7. Mikheenko, A., Prjibelski, A., Saveliev, V., Antipov, D. & Gurevich, A. Versatile genome

assembly evaluation with QUAST-LG. *Bioinformatics* **34**, i142–i150 (2018).
