## Supplementary figures and images for "A granular view of the traits controlling soil bacterial community reconstruction following a prescribed burn"

### Extended Data Figure 1

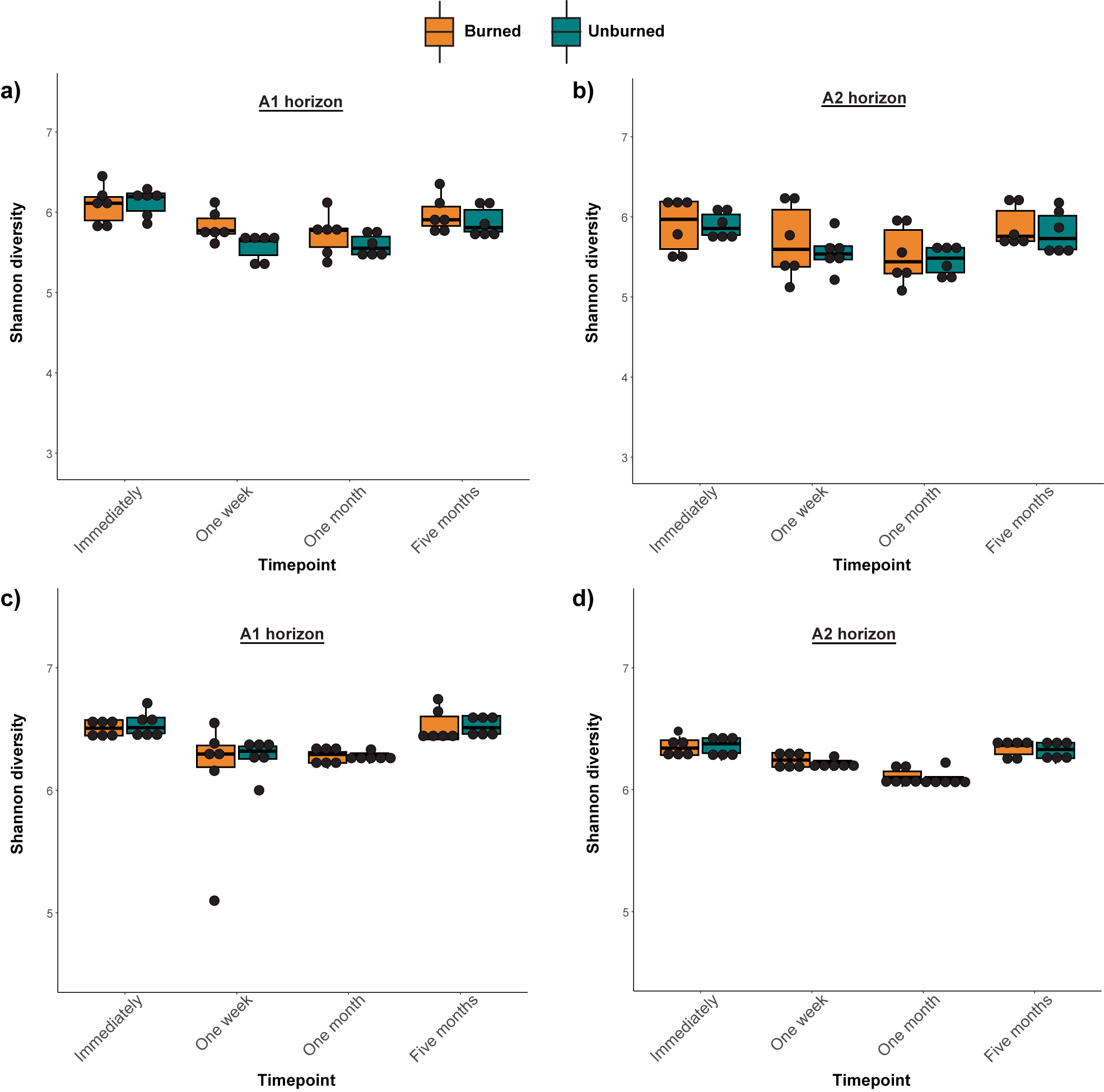

### Extended Data Figure 2

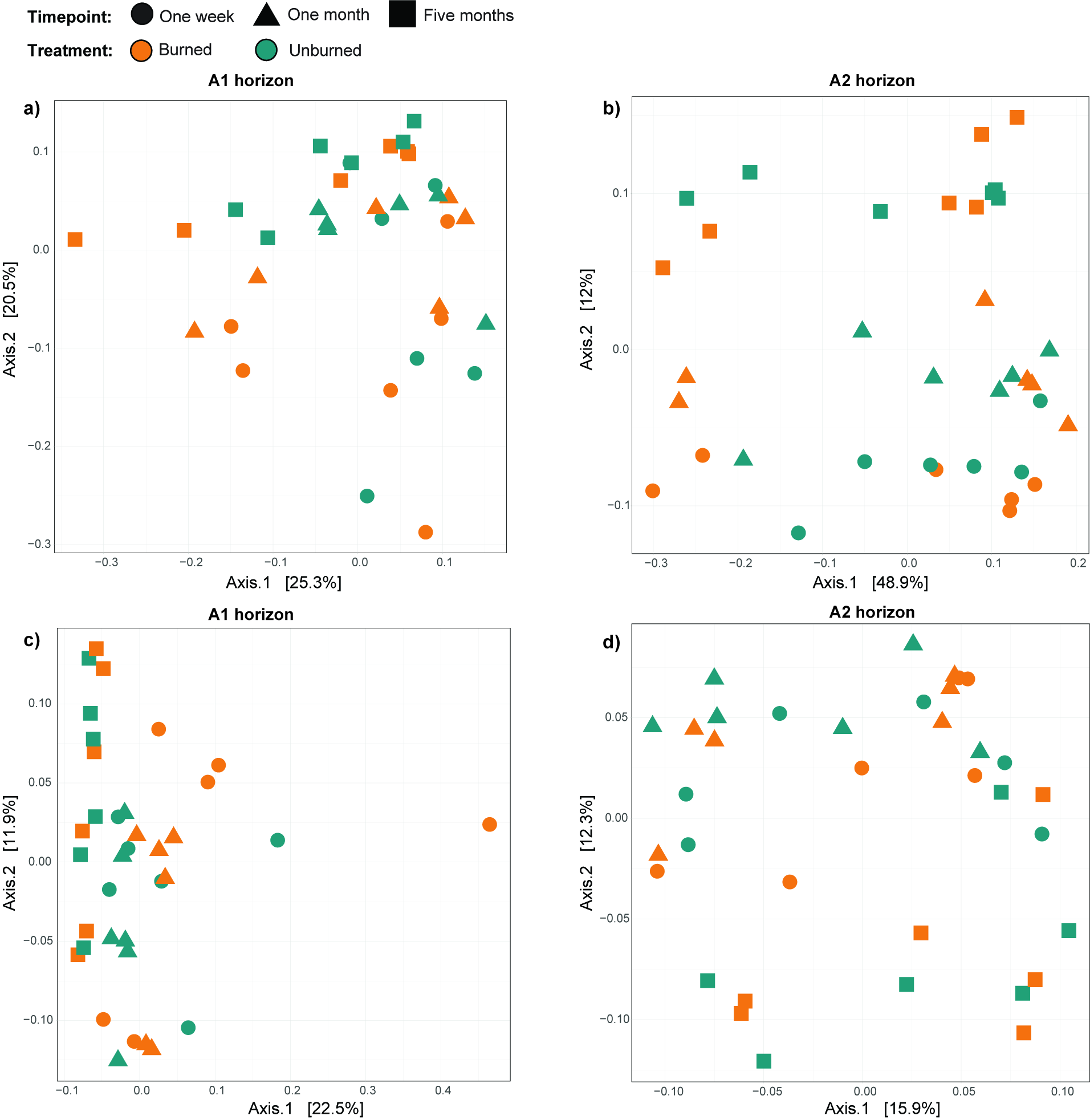

### Extended Data Figure 3

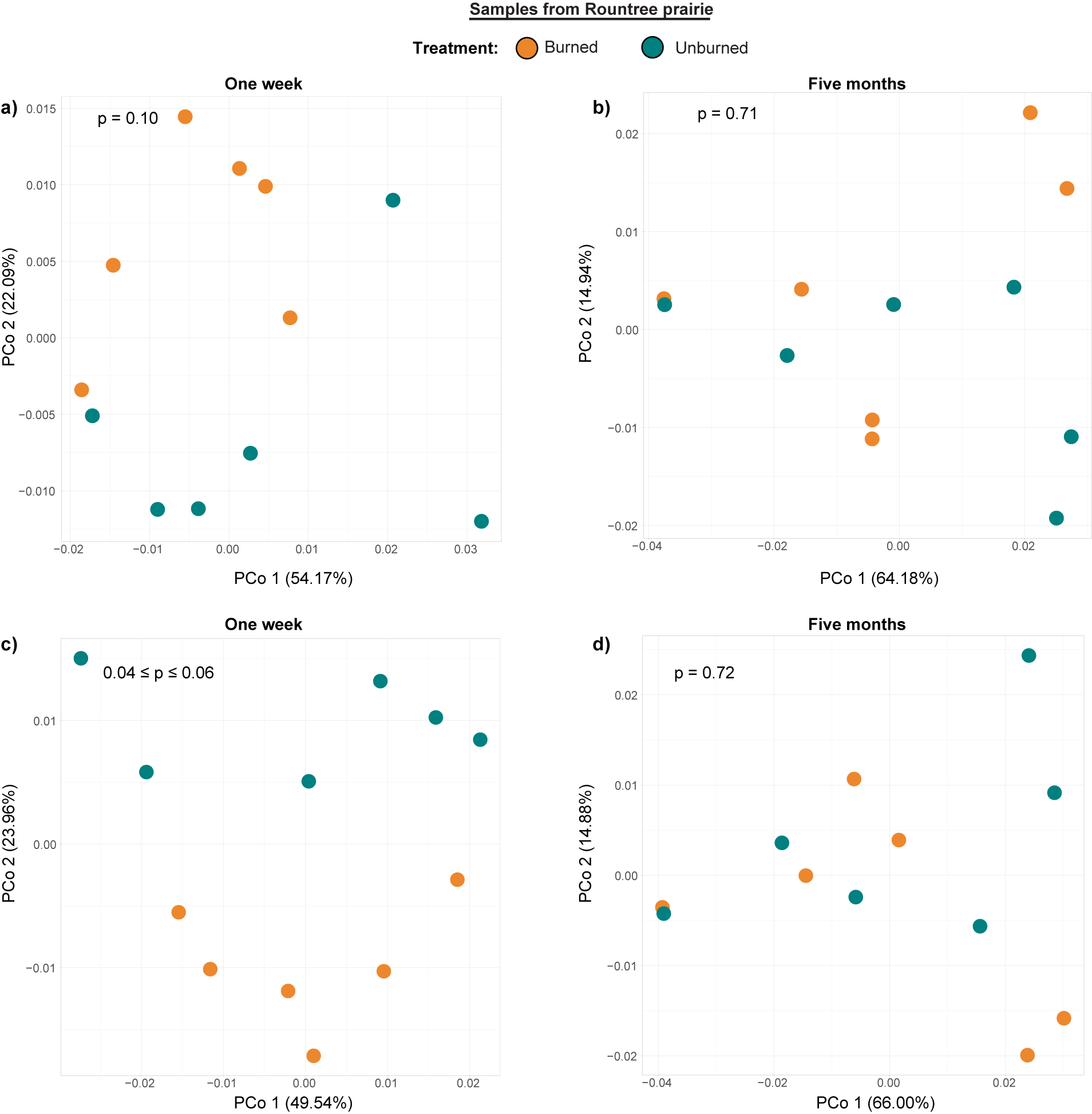

### Extended Data Figure 4

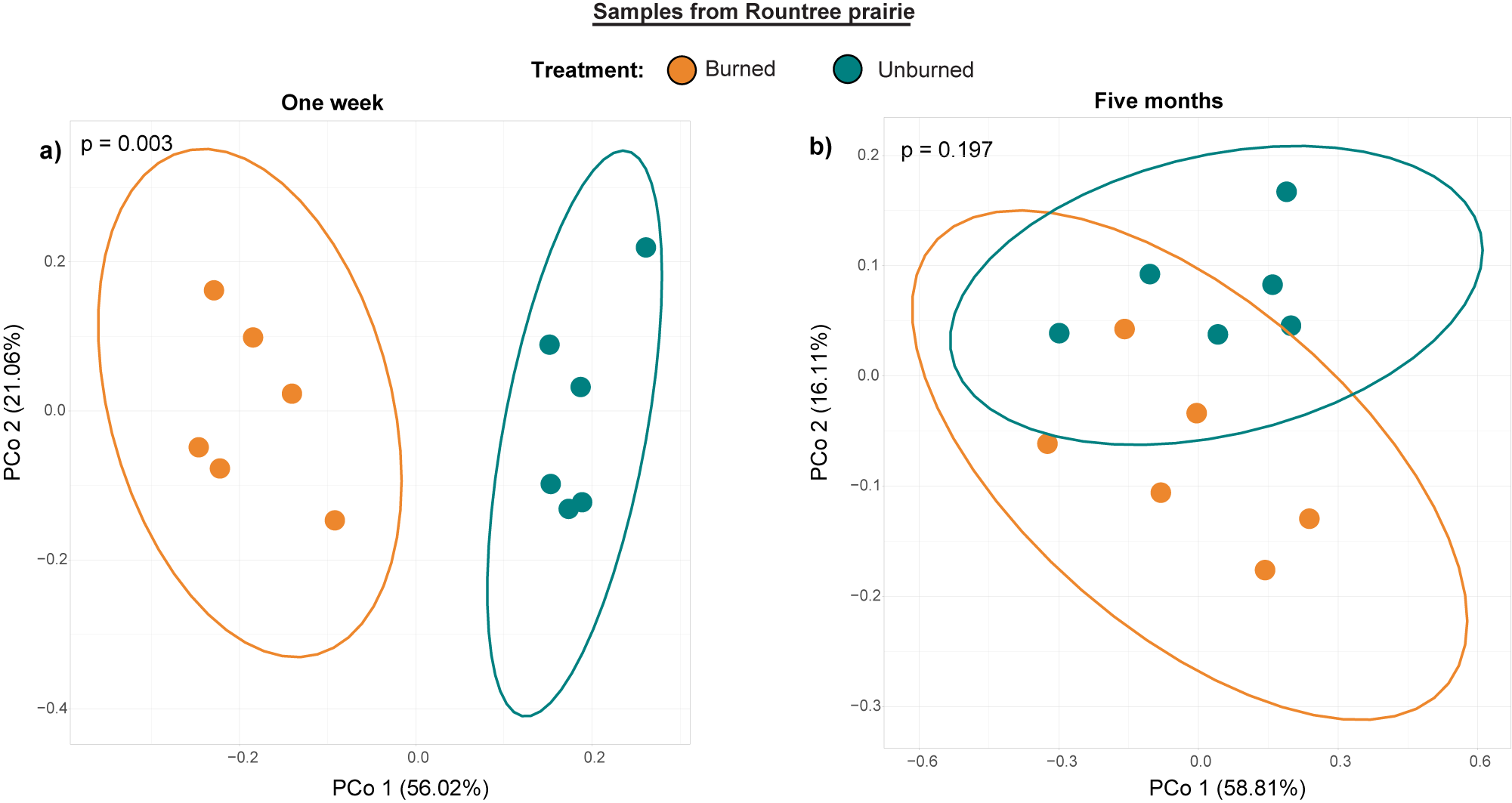

### Extended Data Figure 5

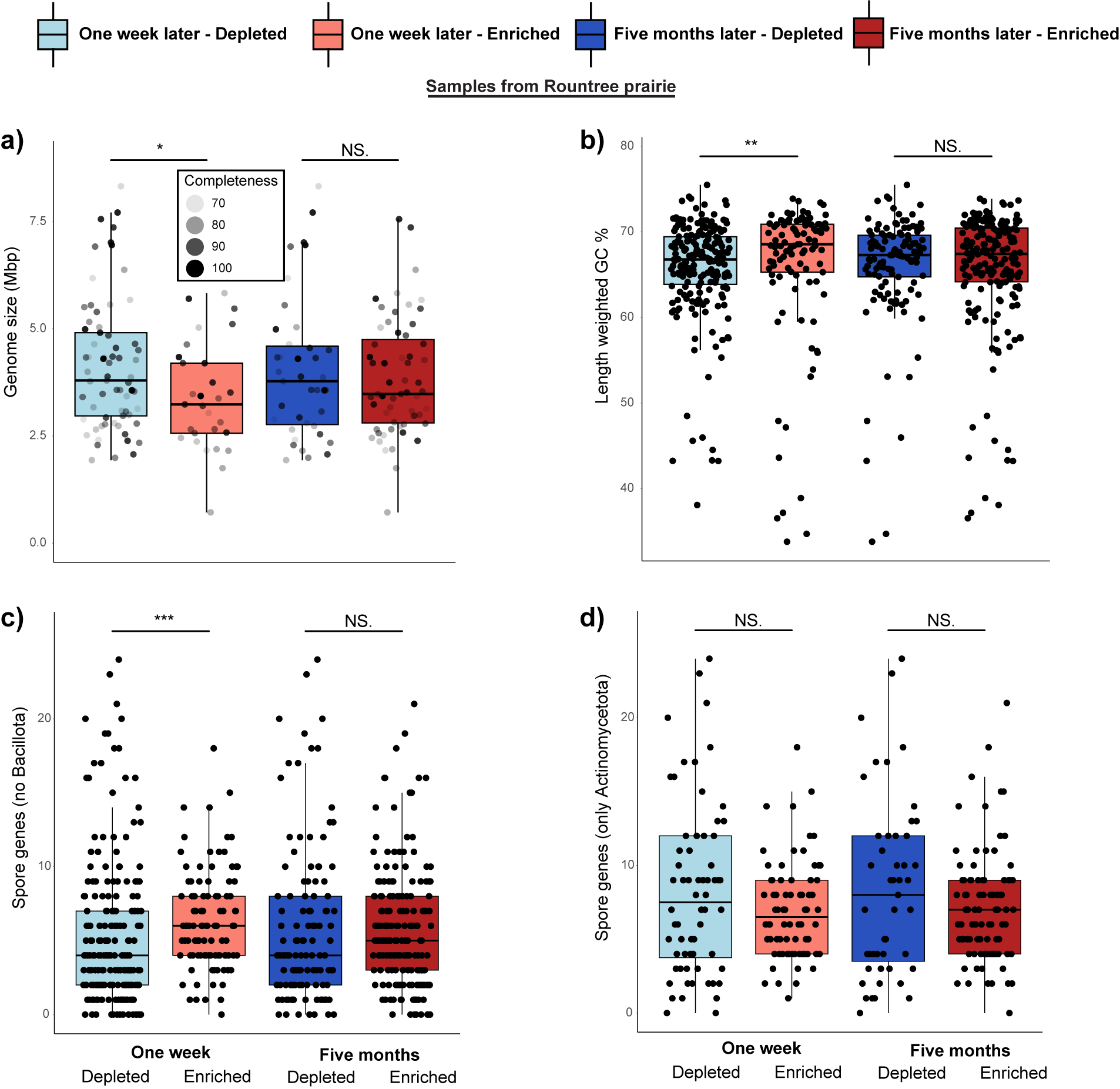

### Extended Data Figure 6

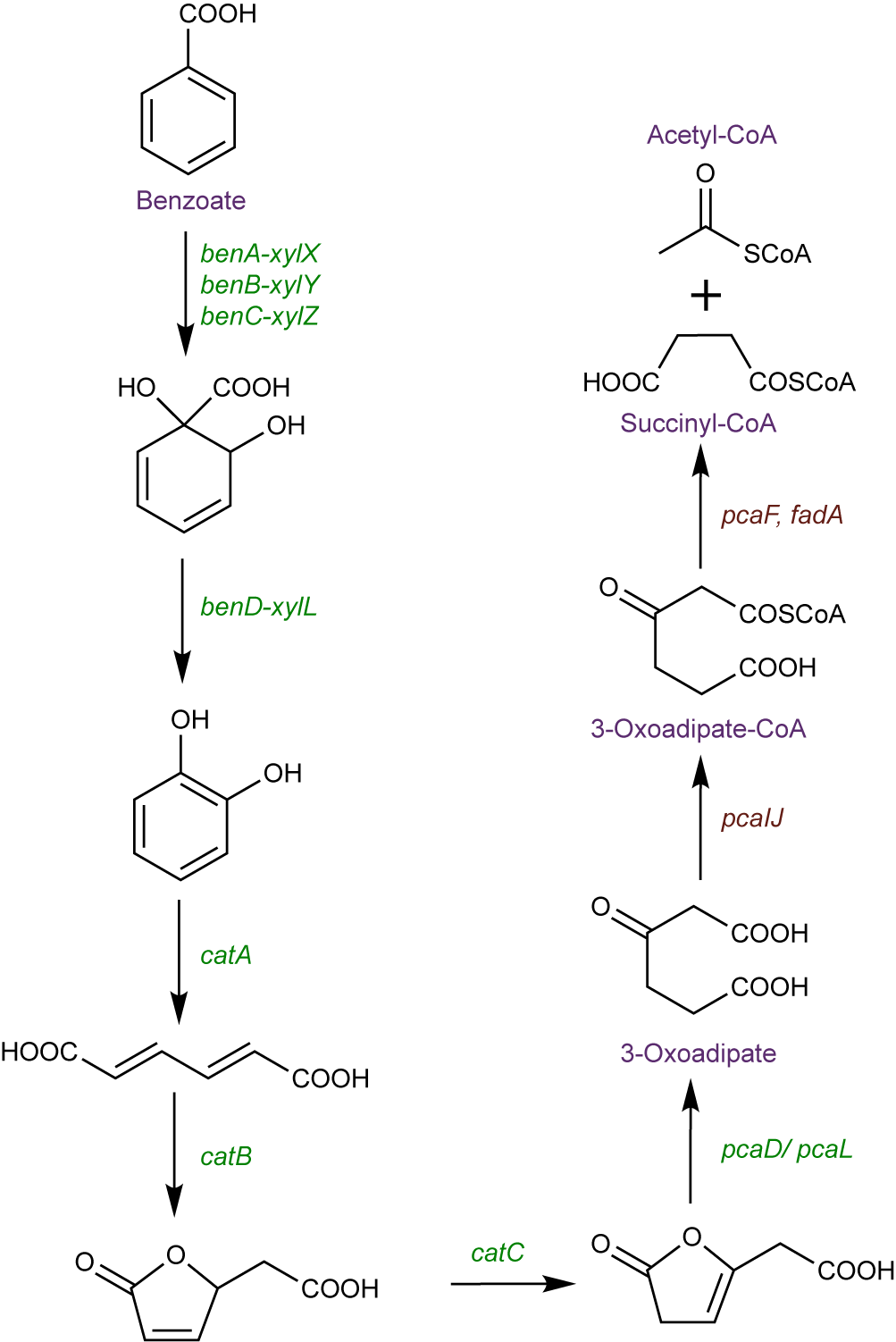

### Extended Data Figure 7

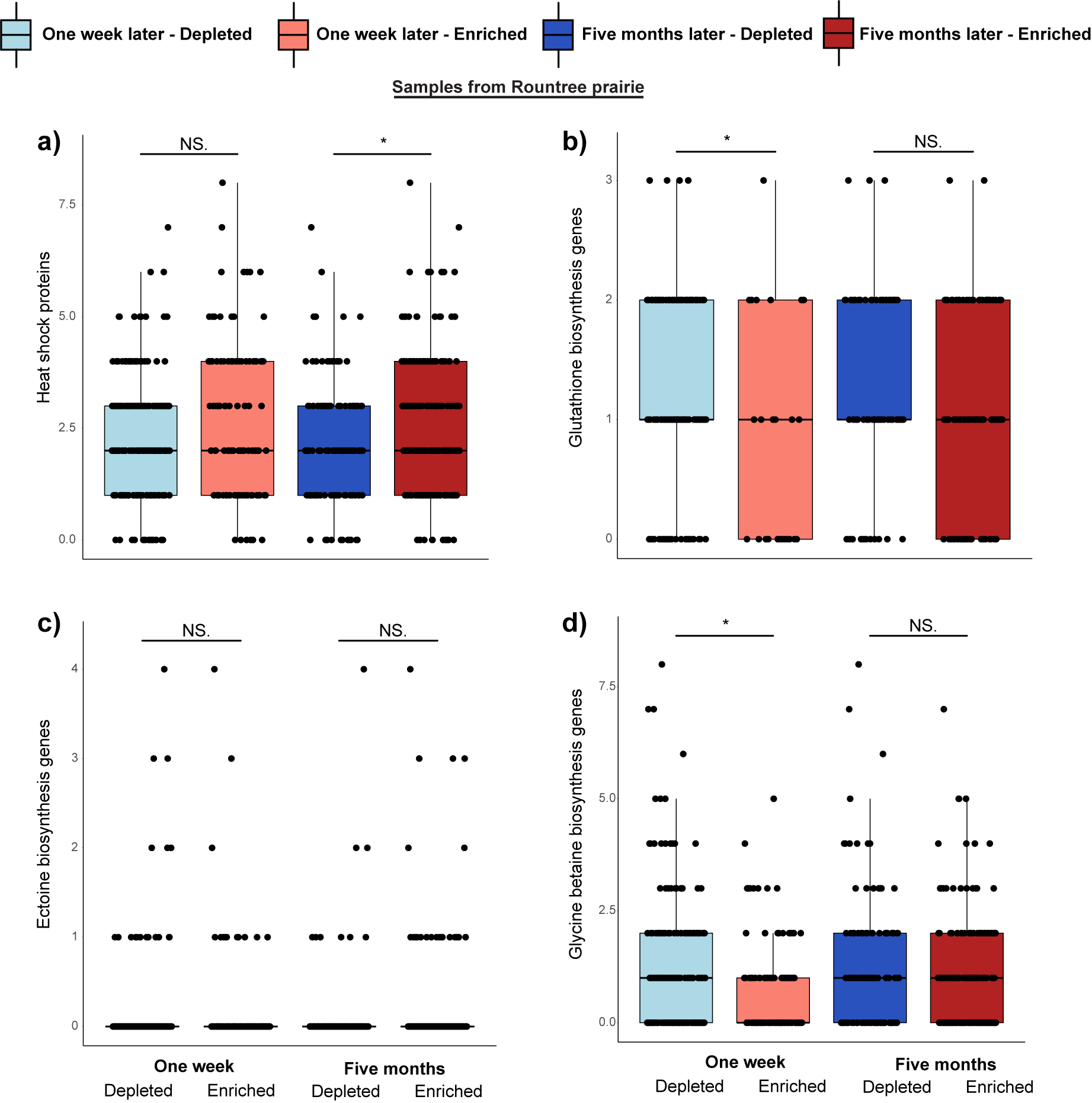

### Extended Data Figure 8

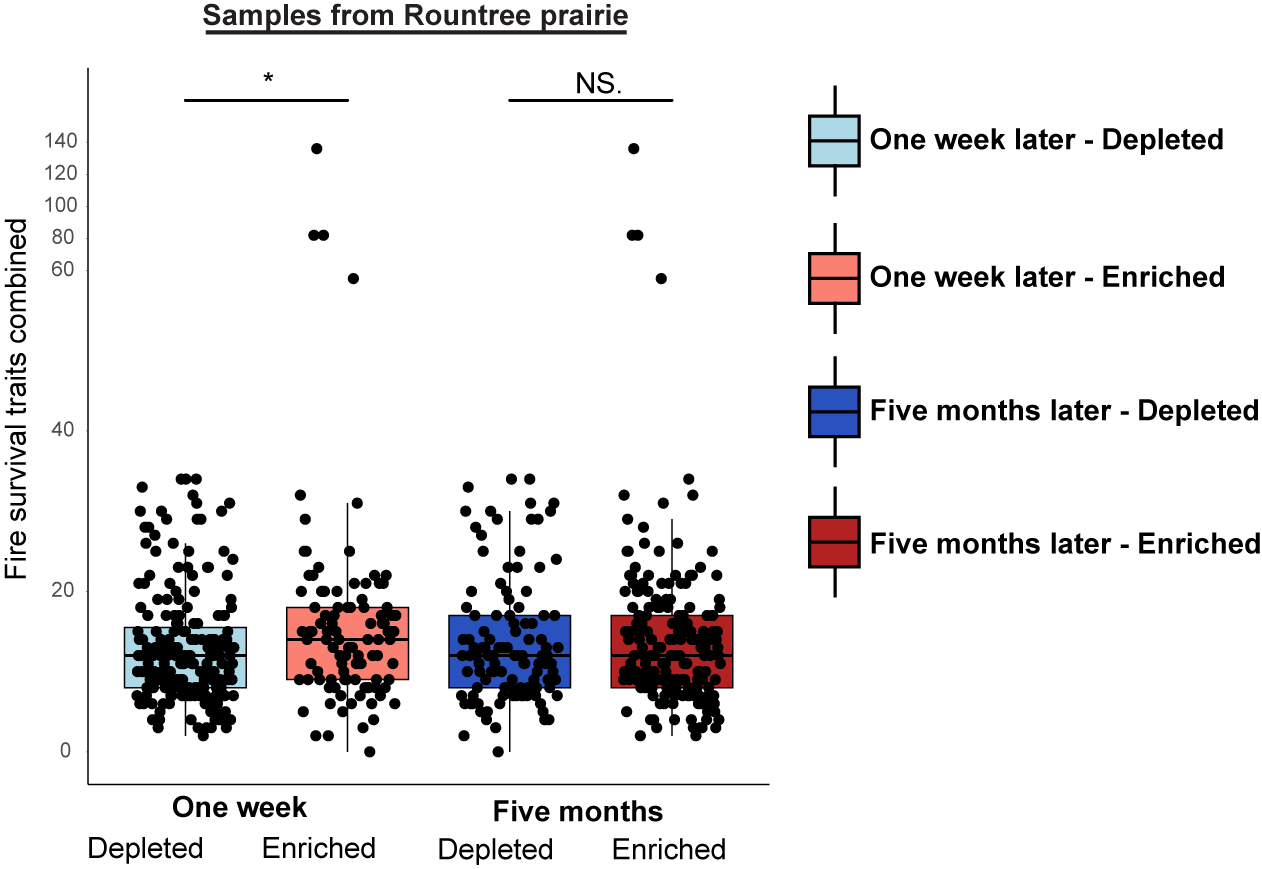
